# Neural stem cell expressed vascular endothelial growth factor maintains the neurogenic vascular niche of the adult mouse hippocampus

**DOI:** 10.1101/2021.08.10.455866

**Authors:** Tyler J. Dause, Jiyeon K. Denninger, Robert Osap, Ashley E. Walters, Joshua D. Rieskamp, Akela A. Kuwahara, Elizabeth D. Kirby

## Abstract

The vasculature is a key component of adult brain neural stem cell (NSC) niches. In the adult mammalian hippocampus, NSCs reside in close contact with a dense capillary network. How this niche is maintained is unclear. We recently found that adult hippocampal NSCs express vascular endothelial growth factor (VEGF), a soluble factor with chemoattractive properties for vascular endothelia. Here, we show that NSC-specific VEGF loss led to dissociation of NSCs from local vasculature but surprisingly found no changes in vascular density. Instead, we found that NSC-derived VEGF supports the motility of NSCs themselves via a cell internal signaling loop. We further found that this internal autocrine loop also independently maintained NSC quiescence cell autonomously. Combined, our findings suggest that NSCs maintain their populations via 2 mechanisms that are dependent on cell internal VEGF signaling: self-stimulated motility towards vasculature and cell autonomous maintenance of quiescence programs.

## Introduction

Neural stem cells (NSCs) and their progeny reside in two primary neurogenic niches in the adult mammalian brain, the subventricular zone (SVZ) and the dentate gyrus (DG) of the hippocampus. Within the adult DG niche, NSCs give rise to intermediate progenitor cells (IPCs) that go on to generate new neurons, a process that is conserved across most land-based mammals studied to date (Charvet and Finlay, 2018; Kempermann, 2016). Evidence in rodent models supports the functionality of this adult neurogenesis, showing that newborn neurons can modulate DG circuit properties (Li et al., 2023; McAvoy and Sahay, 2017; Tuncdemir et al., 2019) and support spatial learning and memory and affective behaviors (Miller and Sahay, 2019). Adult neurogenesis has also generated interest as a potential therapeutic target in a number of diseases and disorders in which hippocampal memory function is impaired (Choi and Tanzi, 2023; McAvoy and Sahay, 2017; Wander and Song, 2021).

The vasculature is a key component of stem cell niches throughout the body (Gómez-Gaviro et al., 2012; He et al., 2014; Verma et al., 2018), including the adult brain niches (Palmer et al., 2000; Shen et al., 2008; Sun et al., 2015; Tavazoie et al., 2008). The adult DG vascular niche is characterized by a dense network of planar capillaries concentrated in the subgranular zone (SGZ) where NSCs and their daughter IPCs (together, NSPCs) reside. Within the SGZ, IPC bodies make close contact with local vessels, using the vessels to guide their migration tangentially through the SGZ away from their parent NSCs before differentiating into a neuronal phenotype (Palmer et al., 2000; Sun et al., 2015). NSCs also achieve unique vascular contact via their radial glia-like process, which extends through the granule cell layer to wrap around vessels in the molecular layer (Licht et al., 2020a; Moss et al., 2016). This vascular niche is hypothesized to provide NSPCs with access to circulating factors as well as signaling from the endothelial cells that comprise vessels, both of which support NSC maintenance and neurogenesis (Kim et al., 2021a; Licht and Keshet, 2015; Villeda et al., 2014; Yousef et al., 2019) Despite wide-spread recognition of the importance of the adult DG vascular niche, there is little known about what signals maintain it.

We previously showed that adult DG NSCs synthesize vascular endothelial growth factor (VEGF) (Kirby et al., 2015), a pleiotropic soluble protein with multiple signaling roles in the adult brain (Lange et al., 2016). We showed that self-produced NSPC-VEGF is necessary for maintenance of the NSPC pool (Kirby et al., 2015), and suggested that this may rely on autocrine signaling via VEGF receptors expressed on NSCs. However, VEGF also can have potent mitogenic and chemoattractant roles for vascular endothelial cells. The role of NSPC-VEGF in DG vascular niche maintenance was left largely unexplored in our previous work.

In this study, we investigated the hypothesis that NSPCs maintain their own vascular niche of the adult mouse DG via production of VEGF. In support of this hypothesis, we found that loss of NSPC-specific VEGF led to disruption of the neurovascular niche. Unexpectedly, however, niche disruption was not accompanied by detectable changes in vessel density, vascular endothelial survival, angiogenesis, or changes to blood-brain-barrier (BBB) proteins. Instead, VEGF loss caused impaired NSC migration, which derived from loss of VEGF intracellular autocrine (intracrine) signaling. We further showed that the reliance of in vitro NSCs on VEGF signaling we have previously reported also relies on an intracrine loop. Overall, our findings suggest that NSPC-derived VEGF supports adult neurogenesis via intracrine signaling that cell autonomously supports both NSPC proximity to vessels and a direct intracellular stemness program simultaneously.

## Results

### Hippocampal NSPCs and astrocytes express VEGF

Our first step to better understand how NSPC-derived VEGF might affect their vasculature niche was to more fully characterize VEGF production by different DG cell types in vivo. First, we quantified the VEGF-expressing populations of the DG using a VEGF-GFP transcriptional reporter mouse (Figure 1A). We found VEGF-GFP expression in the granule cell layer and SGZ primarily in GFAP+/SOX2+ radial glia-like NSCs (RGL-NSCs) and GFAP-/SOX2+ IPCs, while VEGF-GFP expression in the hilus was mostly in GFAP+ astrocytes (Figure 1B,C). Nearly all astrocytes, RGL-NSCs or IPCs were VEGF-GFP+, with little expression in neuronally committed neuroblasts and immature neurons (Figure 1D). To explore relative VEGF expression in a more unbiased manner, we used published single-cell RNA sequencing datasets derived from the adult mouse DG. We found that RGL-NSCs and their IPC progeny expressed *Vegfa* at levels intermediate between two other known producers of VEGF, astrocytes and endothelia (Figure 1E) (Batiuk et al., 2020; Hochgerner et al., 2018; Walker et al., 2020). Analysis of additional published RNAseq datasets revealed that hippocampal NSC *Vegfa* expression increases 1285 ± 45% from embryonic to adult ages (Berg et al., 2019) (Figure 1F) and is 1254 ± 102% greater in hippocampal NSPCs than in NSPCs from the SVZ (Adusumilli et al., 2021) (Figure 1G). To further confirm these sequencing-based results, we performed RNAscope® in situ hybridization in adult mouse DG fixed tissue sections. We used GFAP immunolabeling to identify putative RGL-NSCs and astrocytes based on their unique morphologies. GFAP+ putative RGL-NSCs expressed *Vegfa* but 62.43 ± 4.62% less than putative GFAP+ astrocytes (Figure 1H,I). Taken together, these data further validate our previous findings that NSPCs are a significant source of VEGF in the adult mouse DG, though they produce less *Vegfa* per cell at the RNA-level than astrocytes.

**Figure 1:**
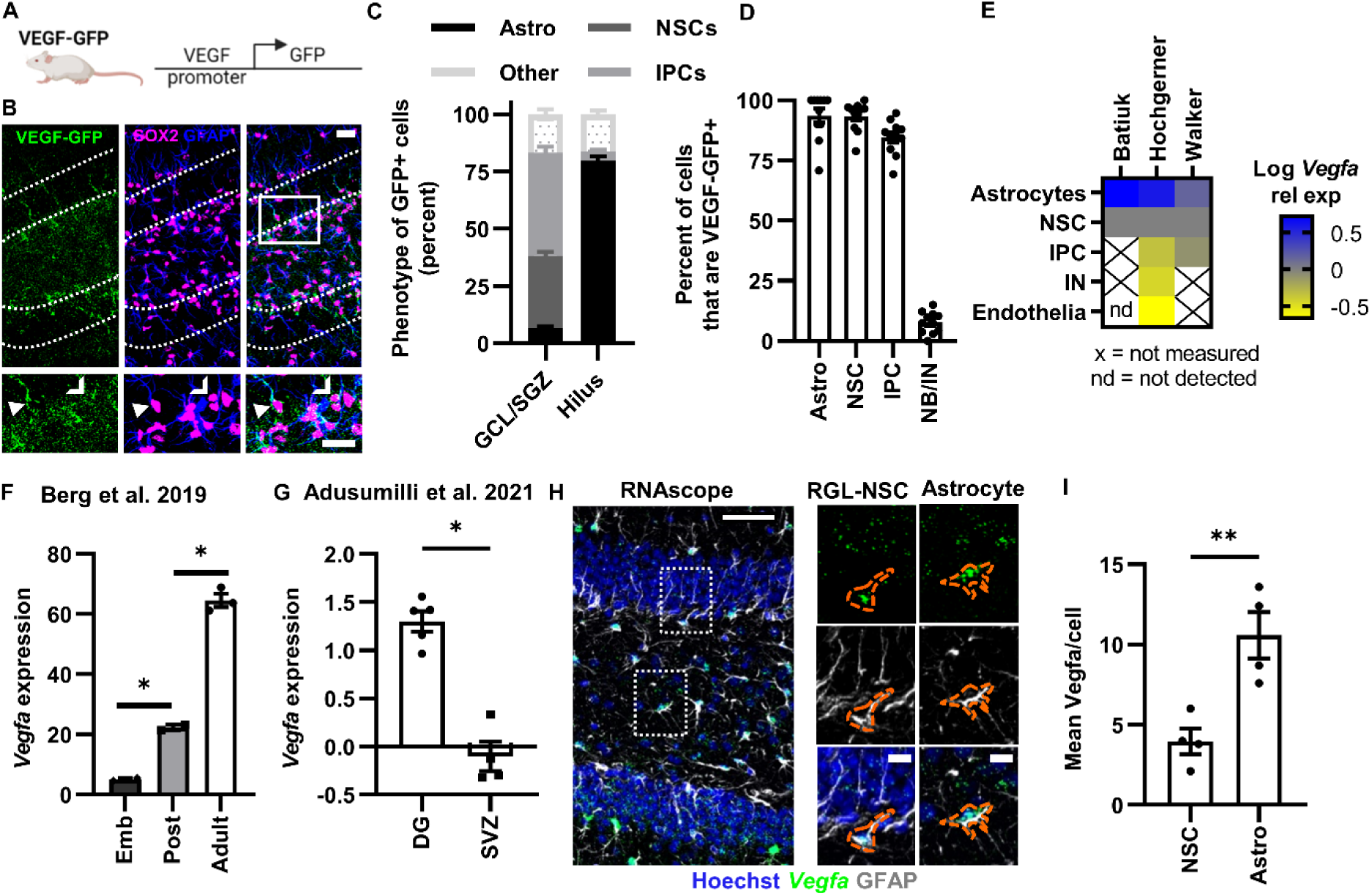
DG NSPCs and astrocytes express VEGF. A) Diagram of VEGF-GFP mouse model. B) Representative immunofluorescent images of GFP, GFAP, and SOX2+ in the DG. Subset images below of GFP+/GFAP+/SOX2+ RGL-NSCs, GFP+/GFAP-/SOX2+ IPCs and GFP+/GFAP+/SOX2+ astrocytes. Arrowhead = GFP+ RGL-NSCs. Chevron = GFP+ İPC. Arrow = GFP+ astrocyte. C) Percent of GFP+ cells of each phenotype: Astrocytes, RGL-NSCs, IPCs and Other. N = 11 mice. D) Percent of Astrocytes, NSCs, IPCs and NB/IN that are GFP+. N = 11 mice. E) Heatmap of *Vegfa* expression from independently published single-cell RNA sequencing datasets (Batiuk et al., 2020; Hochgerner et al., 2018; Walker et al., 2020). IN = immature neurons, nd = not detected. F) *Vegfa* expression across development in Hopx+ NSCs isolated from mouse DG and quantified via bulk RNA sequencing originally published in Berg et al., 2019. N -2-3 mice. G) *Vegfa* expression of NesGFP+ NSPCs in the DG or SVZ quantified via bulk RNA sequencing originally published in Adusumilli et al., 2021. N =4-5 mice. H) *Vegfa* in situ hybridization co-labeled with GFAP antibody in adult mouse DG. Subset images of *Vegfa* in situ hybridization co­labeled with GFAP antibody in adult mouse DG. I) Mean *Vegfa* RNA per cell in RGL-NSCs and astrocytes. N = 4 mice. Mean ± SEM plus individual mice shown. Scale bars represent B,C) 20 µM H) 50 µM, subset 10 µM *p < 0.05; **p < 0.01

### NSPC-VEGF loss disrupts the DG neurogenic vascular niche

To examine the role of NSPC-derived VEGF in maintaining the hallmark features of the DG vascular niche, we used a NSPC-specific, inducible VEGF knockdown model. To knockdown VEGF specifically in NSCs and their progeny, we used NestinCreER^T2+/-^;VEGF^lox/lox^;ROSA-

LoxSTOPLox-EYFP^+/+^ mice. We have previously shown that tamoxifen (TAM) administration in these mice drives recombination specifically in NSPCs and results in loss of about 1/3 of total DG VEGF compared to wildtype littermates (Denninger et al., 2023; Kirby et al., 2015). Here, we treated NestinCreER^T2+/-^;VEGF^lox/lox^;ROSA-LoxSTOPLox-EYFP^+/+^ (iKD) mice and NestinCreER^T2+/-^;VEGF^wt/wt^;ROSA-LoxSTOPLox-EYFP^+/+^ (WT) littermates with TAM to induce recombination and EYFP expression in NSCs, IPCs and their progeny (Figure 2A). Using immunolabeling of fixed brain sections collected 21d after TAM, we quantified the shortest distance between CD31+ vasculature and the cell body center of EYFP+/GFAP+ RGL-NSCs, EYFP+/MCM2+ IPCs and random cells (Figure 2B). We expressed this data as vessel association, which is the measured cell’s distance to the closest CD31+ vessel minus that of the average of random cells in WT mice. In this measure, a cell with an association measure of 0 is equidistant to the vasculature as a random cell. Cells with a score < 0 are closer to vessels than random distribution would predict while cells with a score of > 0 are farther from vessels than random distribution would predict (Figure 2C).

**Figure 2:**
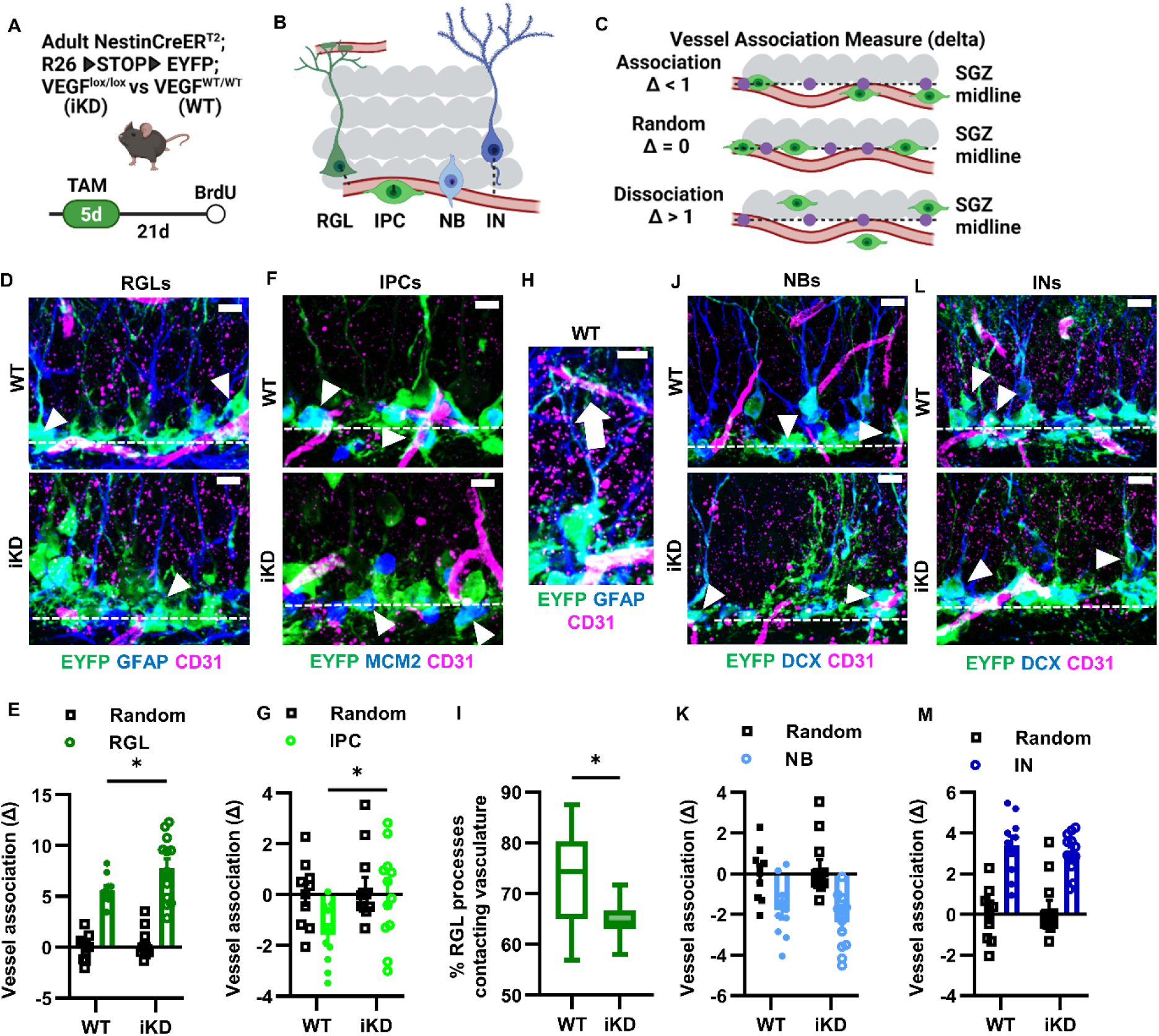
Neurogenic vascular niche disruption after NSPC-VEGF loss. A) Diagram of experimental design and timeline. B) Diagram of vascular distance measurements in NSCs and progeny. C) Diagram explaining vessel association measure. D) Representative immuπofluorescent images of GFAP+EYFP+ RGL-NSCs and CD31+ endothelia 21 d after NSPC-VEGF knockdown. Arrowheads indicate EYFP+ RGL-NSCs. E) Vessel association measurement in WT or İKD RGL-NSCs compared to random points in SGZ. F) Representative immuπofluorescent images of MCM2+EYFP+ IPC and CD31+ endothelia 21d after NSPC-VEGF knockdown. Arrowheads indicate EYFP+ IPCs. G) Vessel association measurement in WT or İKD IPCs compared to random points in SGZ. H) Representative immuπofluorescent image of GFAP+EYFP+ RGL-NSC radial process contacting CD31+ endothelia 21 d after NSPC-VEGF knockdown. Arrow indicates vascular contact. I) Comparison of the percentage of RGL-NSCs whose radial process contacted the vasculature. J) Representative immuπofluorescent images of DCX+EYFP+ neurablasts (NB, horizontal morphology) and CD31+ endothelia 21d after NSPC-VEGF knockdown. Arrowheads indicate EYFP+ neuroblasts. K) Vessel association measurement in WT or İKD neuroblasts compared to random points in SGZ. L) Representative immuπofluorescent images of DCX+EYFP+ immature neurons (IN, dendritic morphology) and CD31+ endothelia 21 d after NSPC-VEGF knockdown. Arrowheads indicate EYFP+ immature neurons. M) Vessel association measurement in WT or İKD immature neurons compared to random point in SGZ. Mean ± SEM plus individual mice shown. N = 10 WT, 12 İKD. Scale bars = 10 µm. *p < 0.05. D,F,J,L) Dashed line indicates lower border of granule cell layer. See also Supplementary Figure 1.

Among WT mice, we found as expected that RGL-NSCs somas were further than random from CD31+ vessels while IPCs were closer to vessels than random distribution (Fig 2D-G). In iKD mice, RGL-NSC and IPC somas were both significantly further from the vasculature compared to WT controls (Figure 2D-G). RGL-NSC somas in iKD mice were 1.4 fold farther from vessels than those in WT mice, with no difference evident for random point distance to vessels. The IPC association with vessels evident in WT mice was completely disrupted in iKD mice, with iKD IPCs showing similar vessel proximity as random cells. We also found that RGL-NSC process contact with vessels was significantly disrupted 21d after TAM (Figure 2H,I). Similar to IPCs as a whole, proliferative cells labeled with the mitotic marker BrdU 2h before perfusion were closer to vessels than random in WT mice and this association was disrupted in VEGF iKD mice (Supplementary Figure 1A,B). These data show significant disruption of vascular proximity of NSPC bodies after loss of NSPC-derived VEGF.

We next examined neuroblast and immature neuron progeny of NSPCs. These cell types do not normally self-synthesize notable quantities of VEGF (Figure 1A), but neuroblasts migrate tangentially away from their parent RGL-NSCs using the vasculature as a scaffolding (Sun et al., 2015). As neuroblasts differentiate into immature neurons, they then migrate radially into the GCL becoming more avascular (Sun et al., 2015). We categorized doublecortin (DCX)+ cells into neuroblasts and immature neurons morphologically, similarly to previously published methods (Plümpe et al., 2006). Similar to previous work (Sun et al., 2015), we found that immature neurons were generally farther from vessels than neuroblasts, which were frequently contacting CD31+ endothelia (Figure 2J-M). There was no difference in vessel association of neuroblasts or immature neurons between WT and iKD mice (Figure 2J-M). All together, these data suggest that NSPC-specific VEGF is necessary to maintain the vascular proximity of NSPCs, with no effect on their neuroblast and immature neuron progeny.

### NSPC-VEGF loss has no detectable effect on DG vasculature

VEGF is a powerful mitogen and chemokine for endothelial cells of the vasculature (Olsson et al., 2006). We therefore next investigated whether loss of NSPC-VEGF may disrupt maintenance of vasculature in the DG cell layers. First, we examined the effect of NSPC-VEGF loss on angiogenesis by colabeling CD31+ endothelia with MCM2, a marker of the cell cycle (Krstulja et al., 2008). In adulthood, brain vasculature is generally considered quiescent (non-proliferative and stable) (Wälchli et al., 2023). Consistent with this non-proliferative nature of adult vasculature, we found no evidence of angiogenesis in the adult mouse DG in both WT and iKD mice (Figure 3A). We also found no evidence of endothelial cell death, as evidenced by a lack of co-labeling for CD31 with activated Casapase-3, a marker of apoptosis (Figure 3B). Finally, we quantified vascular coverage to determine if NSPC-VEGF loss drove endothelial migration out of the SGZ (for example, towards astrocytes in the molecular layer and hilus). Using immunolabeling for CD31+ area, we found that the SGZ and molecular layer were overall more highly vascularized than the granule cell layer and hilus but, there was no difference between WT and iKD mice in CD31+ area in any DG subregion (Figure 3C,D). These findings suggested no broad vascular gain, loss or migration after NSPC-VEGF loss.

**Figure 3:**
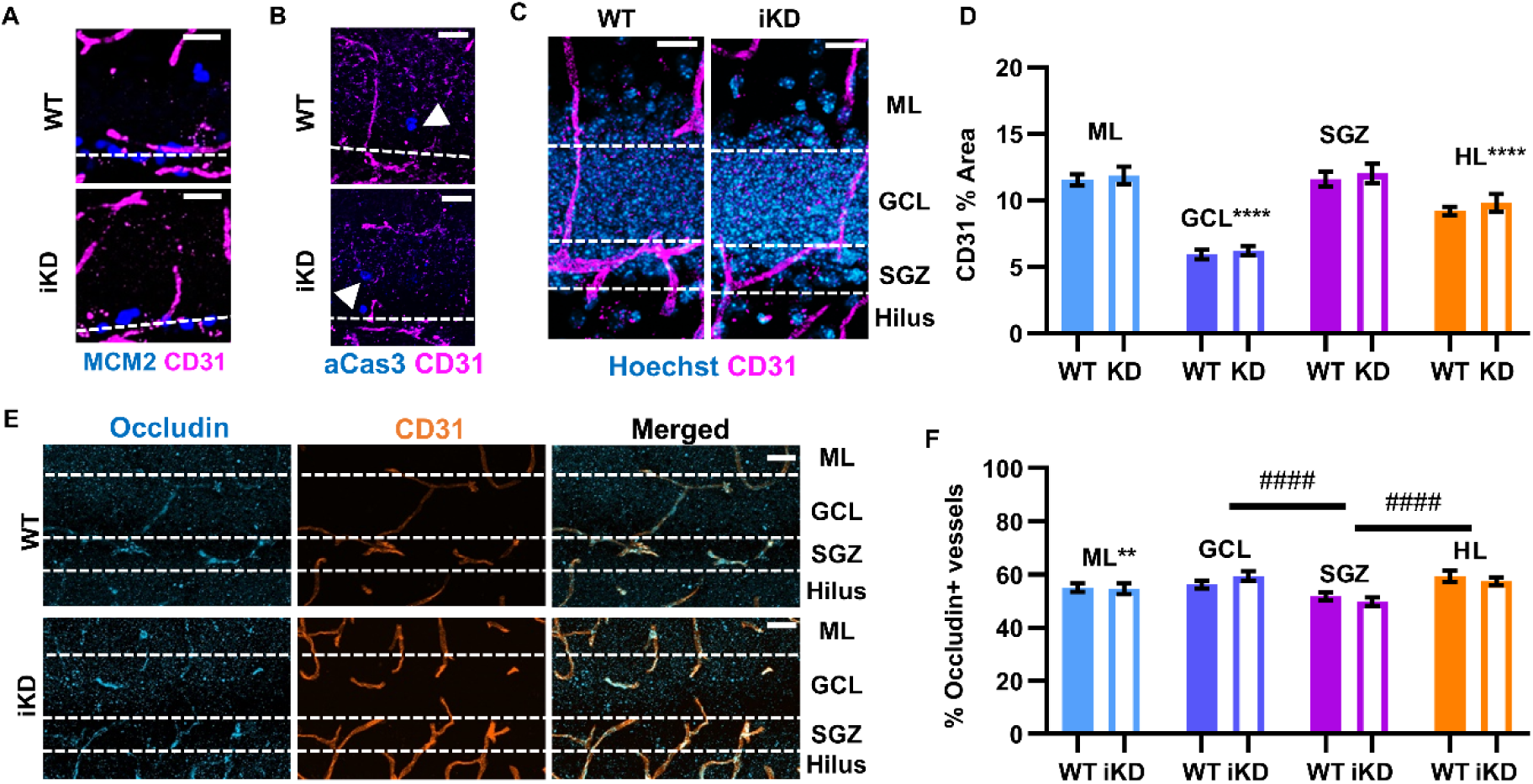
NSPC-VEGF knockdown does not detectably alter the DG vasculature. A) Representative immunofluorescent images of MCM2+ cells and CD31+ endothelia in WT and iKD mice. Dashed lines represents granular cell layer. B) Representative immunofluorescent images of aCas3+ cells and CD31+ endothelia in WT and İKD mice. Arrowheads indicate aCas3+ cell. Dashed lines represents granular cell layer. C) Representative immunofluorescent images of CD31+ endothelia in the DG subregions after NSPC-VEGF knockdown. Hoechst used to stain cell bodies. Dashed lines indicate borders between subregions. D) Comparison of CD31 percent area in the DG subregions in WT and iKD mice 21 d after TAM. E) Representative immunofluorescent images of occludin+/CD31+ endothelia in the DG subregions 21 d after TAM. Dashed lines indicate borders between subregions. F) Comparison of CD31 percent area in the DG subregions in WT and İKD mice 21d after TAM. Mean ± SEM shown. N = 10 WT, 12 iKD mice. Scale bars = 20 µm. **p < 0.01. ****p < 0.0001

In addition to its role in regulating endothelial migration and proliferation, VEGF also stimulates shedding of blood-brain-barrier (BBB) associated proteins on endothelial cells, thereby increasing BBB permeability (Zhang et al., 2000). Occludin is a tight junction membrane protein and a structural component of the BBB that plays an integral role in maintaining BBB impermeability. We therefore used co-labeling of occludin on CD31+ vessels as a measure of BBB integrity (Figure 3E). We found significant, though small, differences between subregions in the percent of occludin co-labeling on CD31+ vessels, with SGZ being the region with the least co-labeling. However, there was no difference in the percentage of occludin+ vessels between WT and iKD mice in any DG subregion (Figure 3F). Taken together, our data were contrary to our expectations and revealed no measurable impact of NSPC-VEGF loss on multiple aspects of adult DG vasculature.

### Loss of NSPC-VEGF disrupts gene expression programs regulating cell migration

Given the apparent lack of change in adult DG vasculature after NSPC-VEGF loss, we next examined changes in NSPCs themselves in more detail. We performed single cell RNA sequencing of RGL-NSCs and their progeny acutely isolated from adult mouse DG of our inducible, NSPC-specific VEGF knockdown model (Figure 4A). 21 days after TAM, when vascular disruption was evident (Figure 2), EYFP+ cells were acutely isolated from dissected DGs using fluorescence activated cell sorting and RNA from single cells was sequenced using a 10X Chromium platform. In total, 5543 WT EYFP+ cells and 5261 iKD EYFP+ cells met inclusion criteria after being captured and sequenced. Uniform manifold approximation and projection for dimension reduction (UMAP) analysis revealed that both WT and iKD cells were present in 11 different subpopulations characterized by gene expression profiles linked to Gene Ontology (GO) terms consistent with cell types across the neurogenic cascade (Figure 4B, Table S1).

**Figure 4:**
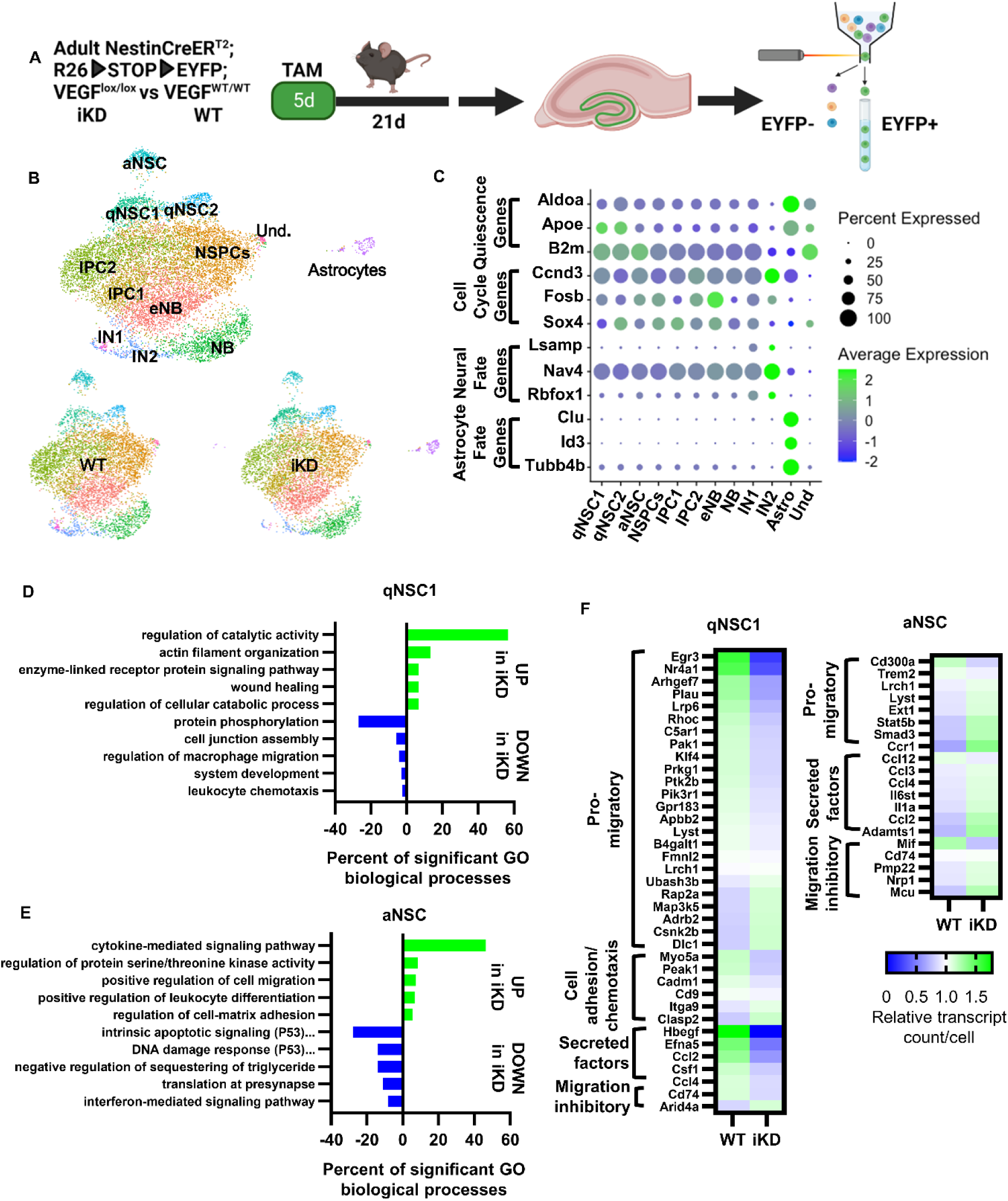
NSPC-VEGF regulates genes related to cell migration in vivo. A) Diagram of experimental design and timeline. B) UMAP of WT and iKD cells yielded 11 subpopulations. Clusters are represented by different colors and phenotypes were assigned by gene expression and GO analysis of DEGs between clusters. C) Dot plot visualization of average expression and percent of cells expressing genes related to quiescence, the cell cycle, neurogenic and gliogenic fate. D) Top 5 GO biological process clusters that were upregulated or downregulated in VEGF iKD qNSC1 cluster. E) Top 5 GO biological process clusters that were upregulated or downregulated in VEGF iKD aNSC cluster. F) Heatmap of upregulated and downregulated genes associated with cell migration in WT and iKD qNSC1 and aNSC clusters. See also Supplementary Figure 2.

To confirm cluster identities, we examined both GO terms associated with cluster marker genes and the expression of known markers of quiescence, the cell cycle, neural fate and gliogenic fate. We identified 3 clusters of NSCs (10.17% of cells), five clusters containing IPCs or NBs (85.09% of cells), two neuronal clusters (3.13% of cells) and one astrocytic cluster (1.24% of cells). The NSC clusters were generally characterized by high expression of quiescence-related genes such as *Aldoa*, *Apoe* and *B2M* and GO terms associated with ribosomal biogenesis, a process previously shown to be high in NSCs (Harris et al., 2021; Shin et al., 2015) (Figure 4C). One of the NSC clusters was characterized by GO terms associated with G1/S transition and cell cycle activation, leading us to define these as active NSCs (aNSCs) (Figure 4C). The IPC and NB clusters showed progressive loss of quiescence gene expression coupled with expression of cell cycle related genes such as *Ccnd3*, *Fosb* and *Sox4* (Figure 4C). GO terms in these populations also highlighted cell cycle-related processes such as regulation of G1/S transition of mitotic cell cycle. We defined one population as a putative mix of NSCs and IPCs (NSPCs) because it showed gene expression and GO terms intermediate between the aNSC and IPC clusters (Figure 4C). The 2 IN populations showed strong expression of neuronal fate genes and GO terms associated with neuronal differentiation. The astrocyte cluster had high expression of quiescence genes coupled with high expression of genes associated with astrocytic fate (*Clu*, *Id3*, *Tubb4b*) and a notable absence of neuronal phenotype and cell cycle genes that were present in NSC, IPC, NB and IN clusters (Figure 4C).

We next identified differentially expressed genes (DEGs) between WT and iKD cells within clusters. Among NSC-containing clusters, qNSC1, qNSC2 and aNSCs showed a high number of DEGs between groups with 210, 1314 and 332 genes respectively. NSPCs, IPC1 and IPC2 clusters, in contrast, only yielded 20, 20 and 72 DEGs respectively. We therefore focused on the qNSC and aNSC clusters. To understand the potential functional significance of these DEGs, we used GO analysis of upregulated and downregulated DEGs coupled with kappa score iterative grouping with ClueGO to collapse similar GO terms. qNSC2 GO terms were primarily associated with changes in differentiation processes, chromosomal regulation, DNA biosynthesis and kinase activities (Supplementary Figure 2A, Table S1), all of which are consistent with our previous finding that loss of NSPC-VEGF induces NSC activation from quiescence and subsequent exhaustion (Kirby et al., 2015).

More interestingly, some of the most frequent GO biological process and molecular function terms for DEGs in iKD qNSC1 and aNSCs compared to WT were associated with cell migration (Figure 4D,E), such as regulation of cell migration, wound healing, leukocyte chemotaxis, and macrophage migration. To further understand the nature of these migration-related changes, we grouped motility-associated up and down DEGs into subcategories based on whether they are implicated in regulating motility of the cell expressing them (pro-migratory, cell adhesion/chemotaxis and migratory inhibitory) or regulating the migration of other cells (secreted factors or their production). We found the majority of DEGs were genes that would be associated with the motility of the NSCs themselves, rather than eliciting migration of other cells (Fig 4F). In qNSCs, we found a preponderance of pro-migratory and chemotactic genes being downregulated in VEGF iKD mice. In aNSCs, in contrast, we observed upregulation of genes associated with both migration promoting and migration inhibiting functions in VEGF iKD mice. Taken together, our single-cell data suggested that loss of NSC-specific VEGF may disrupt NSC migration.

### NSC-VEGF loss inhibits NSC motility

We next sought to validate our transcriptomic results, specifically to confirm that NSC motility relies on self-synthesized VEGF. Rather than focus on individual DEGs, which previous research shows is highly subject to the bias of single cell sequencing for detecting highly expressed genes with low fold changes (Denninger et al., 2022; Squair et al., 2021), we took a functional validation approach using a scratch assay to examine NSC migration in vitro. We have previously established through immunolabeling and bulk and single-cell RNA sequencing that adult DG NSC cultures are composed of primarily quiescent and cycling multipotent NSCs which self-renew when maintained in proliferative conditions (Denninger et al., 2020, 2022). We and others have also previously reported that NSCs in vitro and in vivo express VEGFR2, but not the other major VEGF receptor, VEGFR1 (Cao et al., 2004; Fabel et al., 2003a; Kirby et al., 2015; Wittko et al., 2009). To block autocrine VEGF signaling, we therefore treated adult DG NSCs grown in standard monolayer conditions with a cell permeable inhibitor of VEGFR2 receptor tyrosine kinase activity, SU5416, before making a scratch through the monolayer (Figure 5A). We found that NSC ingression into the scratch was slowed in SU5416 treated NSCs compared to control NSCs (Figure 5B,C).

**Figure 5:**
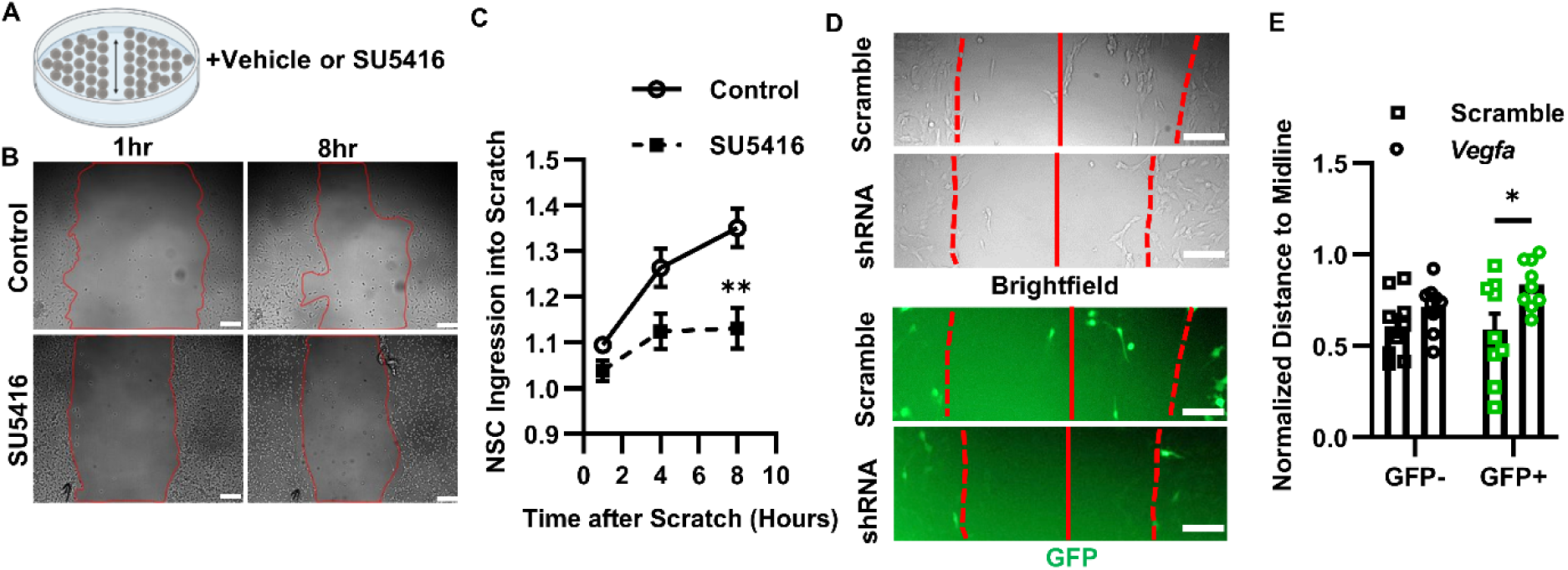
NSC-VEGF regulates cell migration cell autonomously in vitro. A) Diagram of scratch assay experimental design and timeline. B) Representative bright field images of NSC migration into scratch following SU5416 treatment or control after 1 and 8 hrs. Red line = scratch border. C) Comparison of NSC ingression into scratch. Data normalized to initial scratch size. N = 3 wells/experiment, 3 experiments. Mean ± SEM shown. D) Representative images of total (brightfield) and GFP+ NSC migration into scratch following treatment with *Vegfa* shRNA or control virus after 8 hrs. Solid red line = scratch midline. Dotted red line = scratch border. E) Comparison of GFP+ and GFP-cell distance to the midline normalized to 1 hr scratch border distance to midline. N = 3 wells/experiment, 3 experiments. Mean ± SEM and individual wells shown. Scale bars = 100 µm. *p < 0.05. See also Supplementary Figure 2.

To further confirm the necessity of VEGF signaling for NSC migration, we infected NSCs with lentiviral vectors expressing GFP with either a shRNA against *Vegfa* or a scramble shRNA control (Figure 5D). In a separate assay, we first confirmed that *Vegfa* shRNA led to a 60 ± 11% suppression of VEGF compared to scramble shRNA (Supplementary Figure 2B,C). As with VEGFR2 inhibition, ingression into the scratch was impaired by *Vegfa* shRNA expression after 8 hours. Unexpectedly, though, only migration of GFP+, but not GFP-cells, was impaired following VEGF knockdown, suggesting that loss of VEGF may impact NSC migration in a cell autonomous manner (Figure 5D,E). Taken together, our in vitro results support our in vivo transcriptomic results, suggesting that NSC motility relies on self-derived VEGF signaling. They also further suggested an unanticipated cell-autonomous nature of the VEGF signaling that appeared incongruous with the secreted, soluble nature of VEGF.

### NSC VEGF signals through VEGFR2 cell internally

The cell autonomous effects of VEGF knockdown on DG NSC motility suggests that individual NSCs are selectively sensitive to their own VEGF but insensitive to VEGF produced by neighboring cells. This kind of signaling is frequently referred to as intracrine, wherein a ligand signals intracellularly within the cell that synthesizes it, without the need for secretion and binding to extracellularly-facing receptors. A requirement for intracrine signaling is that the cells express both the ligand and receptor simultaneously. We re-examined the datasets from Berg et al., 2019 and Adusumilli et al., 2021, and re-confirmed that hippocampal NSCs express *Kdr* (the transcript for VEGFR2) despite a decline in expression with age (Figure 6A), and that *Kdr* expression is 524 ± 23% higher in DG than in SVZ NSPCs (Figure 6B). We also directly confirmed that almost all (92.5%) of putative RGL-NSCs co-expressed both *Vegfa* and *Kdr* within the same cell using RNAscope in situ hybridization (Figure 6C,D).

**Figure 6:**
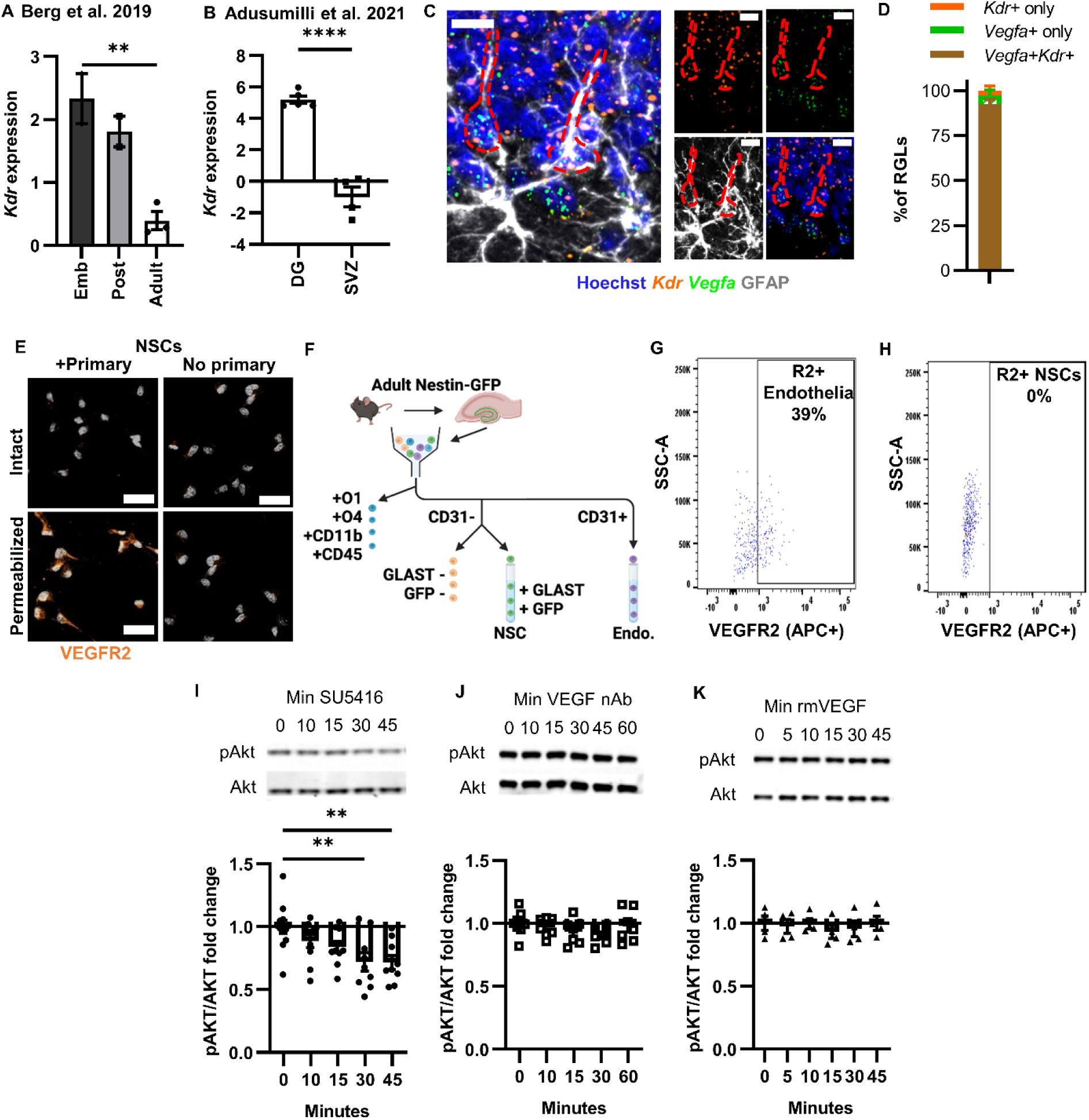
NSCs express cell internal, but not cell surface, VEGFR2. A) *Kdr* expression across development in Hopx+ NSCs isolated from mouse DG and quantified via bulk RNA sequencing originally published in Berg et al., 2019. N = 2-3 replicates; mean ± SEM. B) *Kdr* expression of NesGFP+ NSPCs in the DG or SVZ quantified via bulk RNA sequencing originally published in Adusumilli et al., 2021. N = 4-5 replicates; mean ± SEM. C) *Kdr* and *Vegfa* in situ hybridization co-labeled with GFAP antibody in adult mouse DG. D) Percent of RGL-NSCs that are *Kdr+, Vegfa+* or *Kdr+/Vegfa+.* N = 4 mice/grp; mean ± SEM. E) VEGFR2 immunoreactivity of permeabilized and non-permeabilized cultured DG NSCs in VEGFR2 primary and no primary conditions. F) Diagram of flow cytometry design. G,H) Flow cytometry plot of VEGFR2+ (APC+) labeling in endothelial cells and NSCs. I) Representative western blot of Akt phosphorylation in cultured NSCs after SU5416 over time (top). Fold change in Akt phosphorylation after SU5416 over time (bottom). N = 1-3/grp/exp, 3 exps; mean ± SEM. J) Representative western blots of Akt phosphorylation in cultured NSCs after VEGF nAb over time (top). Fold change in Akt phosphorylation after VEGF nAb over time (bottom). N = 1-3/grp/exp, 3 exps; mean ± SEM. K) Representative western blot of Akt phosphorylation in cultured NSCs after recombinant mouse (rm)VEGF treatment overtime. Fold change in Akt phosphorylation after rmVEGF over time. N = 3/grp/exp, 2 exps; mean ± SEM. See also, supplementary Figure 3.

One reason intracrine signaling can dominate in a cell that expresses both ligand and receptor is the absence of the receptor on the cell surface. We therefore next investigated whether VEGFR2 may not be at the cell surface of cultured NSCs as would be expected. Using immunocytochemical staining of non-permeabilized (intact) and permeabilized cultured DG NSCs, we confirmed that only permeabilized NSCs showed immunoreactivity with an antibody targeting the N-terminus of VEGFR2, suggesting that the N-terminal VEGF binding sites of VEGFR2 are present only within the intracellular compartment of cultured NSCs (Figure 6E). In contrast to NSCs, cultured mouse brain endothelial cells (bEnd.3), which are well established to respond to extracellular VEGF, showed N-terminal VEGFR2-immunoreactivity both on the cell surface and within the intracellular compartment (Supplementary Figure 2A). CRISPRi suppression of *Kdr* expression (αVEGFR2) significantly reduced both intracellular and extracellular VEGFR2 immunoreactivity in b.End3 cells and intracellular VEGFR2 immunoreactivity in NSCs compared to a non-target single guide RNA (NT) (Supplementary Figure 2B-E), confirming the specificity of the VEGFR2 antibody.

To examine VEGFR2 cell surface expression in vivo, we used VEGFR2 immunolabeling and flow cytometry in acutely dissected DG cell suspensions from adult NestinGFP mice (Figure 6F). Based on our published protocol (Denninger et al., 2022), RGL-NSCs can be identified in these mice as O1^-^O4^-^CD11b^-^CD45^-^CD31^-^GLAST^hi^GFP+ cells (Supplementary Figure 2F). We also examined CD31+ endothelia as a positive control where VEGFR2 cell surface labeling should be evident. We found that VEGFR2 was readily detectable on the cell surface of endothelia (31.80% +/-4.70% R2+) but was not detectable on the surface of NSCs (0.14% +/-0.06% R2+) (Figure 6G,H). When we attempted similar immunolabeling as above in permeabilized cells, antibody labeling was not present in either NSCs or endothelial cells, suggesting that antibodies were not effective in these conditions. These findings echo those of others who have shown that VEGFR2 antibodies to be unreliable in permeabilized tissue sections (Licht et al., 2016). Taken together, these findings suggest that NSC express VEGF and VEGFR2 but do not localize VEGFR2 on the cell surface both in vitro and in vivo.

To further confirm a lack of functional VEGFR2 on the cell surface of NSCs, we examined phosphorylation of a known signaling pathway downstream of VEGFR2, PI3K/Akt, in cultured NSCs. Treatment with the cell permeable VEGFR2 inhibitor SU5416 or a separate cell-permeable inhibitor, SU1498, both significantly inhibited phosphorylation of Akt (Figure 6I, Supplementary Figure 3G). Consistent with an intracrine signaling mechanism, VEGF neutralizing antibody (nAb) that specifically binds extracellular VEGF did not alter pAkt (Figure 6J, Supplementary Figure 3H). Similarly, adding exogenous recombinant mouse VEGF to the extracellular media did not alter Akt phosphorylation (Figure 6K, Supplementary Figure 3I). To confirm the biological activity of our reagents, we performed similar experiments in human umbilical vein endothelial cells (HUVECs), which are sensitive to both extracellular and intracrine stimulation of VEGFR2. Both SU5416 and VEGF nAb blocked recombinant mouse VEGF-induced increases in pAkt in HUVECs, confirming that these reagents are bioactive and capable of preventing VEGF signaling through the PI3K/Akt pathway (Supplementary Figure 3J,K). Together, these data further support an intracrine signaling mechanism where NSC-derived VEGF signals exclusively via cell internal VEGFR2.

### NSCs-express sheddases which cleave VEGFR2 from the cell surface

One common cause of insensitivity to extracellular soluble signals is cleavage of cell surface receptors after insertion into the membrane by extracellular sheddases. Sheddases are membrane-bound enzymes that cleave trans-membrane proteins, consisting of members from the ADAM, BACE and MMP protein families. Analysis of our own previously published single-cell RNA sequencing and liquid chromatography tandem mass spectrometry data from cultured adult DG NSCs revealed that multiple sheddases are detectable throughout the cell cycle and in quiescent NSCs, particularly *Mmp15*, *Adam9*, *Adam10* and *Adam12* (Figure 7A,B) (Denninger et al., 2020). Similarly, we analyzed the single-cell RNA sequencing from our in vivo DG NSC-containing populations (Figure 4) and found the expression of multiple sheddases, including *Adam10* and *17* (Figure 7C). Treating cultured NSCs with tumor necrosis factor-α protease inhibitor 1 (TAPI-1), a broad-spectrum inhibitor of sheddases, significantly increased VEGFR2 immunoreactivity on the cell surface of cultured NSCs (Figure 7D,E). TAPI-1 treatment also restored pAkt response to exogenous recombinant VEGF (Figure 7F). Together these data suggest that sheddase cleavage of VEGFR2 from the NSC cell surface inhibits the NSC response to extracellular VEGF.

**Figure 7:**
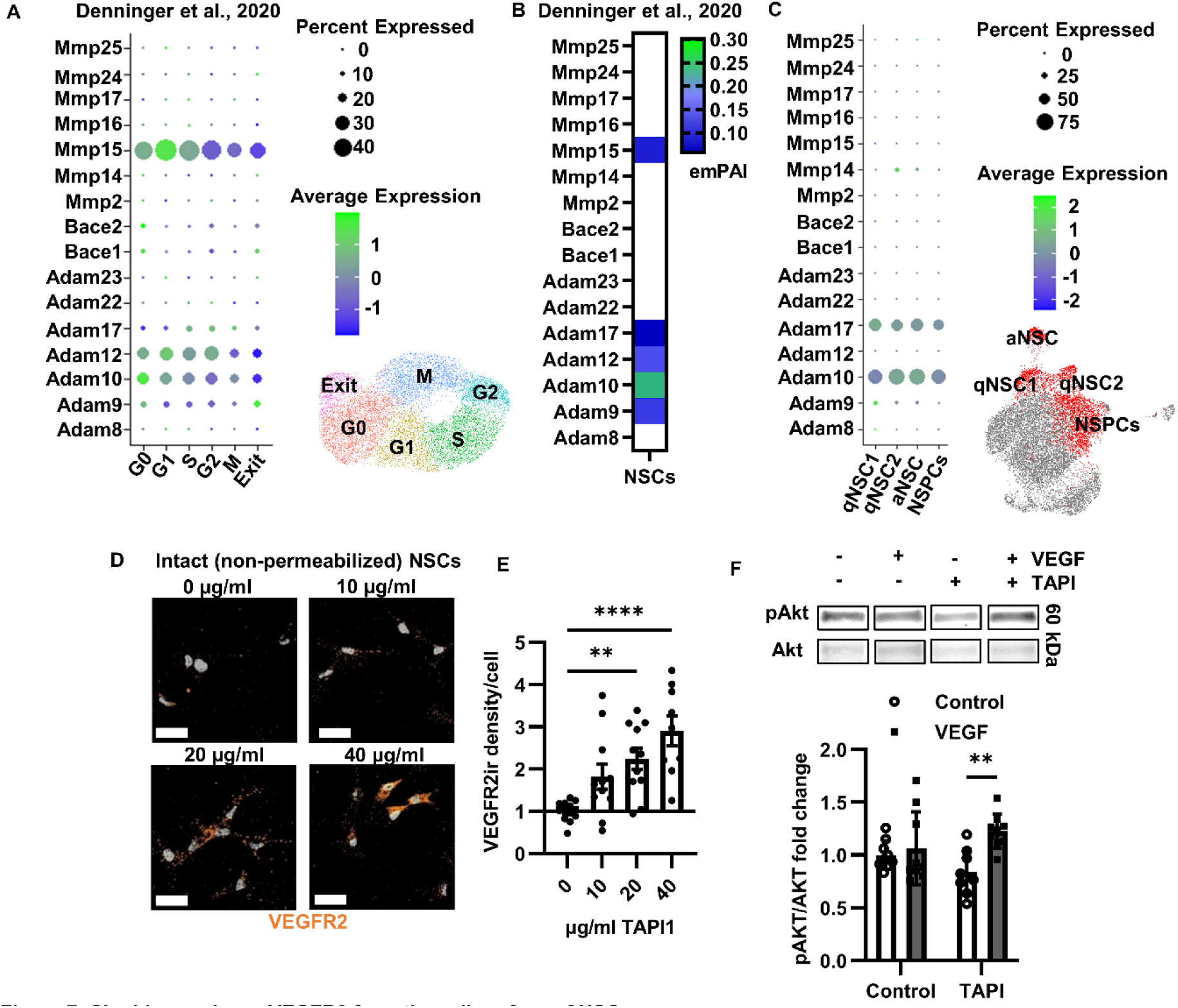
Sheddases cleave VEGFR2 from the cell surface of NSCs. A,B) Sheddase RNA counts from scRNAseq (A) and protein expression (emPAI) from LC-MS/MS (B) of cultured DG NSCs in Denninger et al., 2020. C) Sheddase RNA expression from scRNAseq data in Figure 4, derived from acutely isolated adult hippocampal NSCs. D) Representative VEGFR2 immunoreactivity in intact cultured NSCs following TAPI-1 treatment. Mean cell surface VEGFR2 immunoreactivity per cell following TAPI-1 treatment in cultured NSCs. N = 2-3 wells/exp, 4 exps; mean ±SEM. F) Representative western blots of Akt phosphorylation in cultured NSCs after TAPI-1 and/or recombinant mouse VEGF treatment (top). Fold change in Akt phosphorylation after TAPI and/or recombinant VEGF treatment (bottom). N = 3/grp/exp, 3 exps; mean ± SEM and individual wells shown. Scale bars = 20 µM. **p < 0.01. ****p < 0.0001

### NSPCs rely on cell autonomous VEGF to maintain the neurogenic niche in vivo

Our findings thus far indicated that VEGF signals cell autonomously to support NSC motility in vitro. We next set out to determine if cell autonomous VEGF signaling is also critical in NSPCs in vivo. Our VEGF iKD model (Figure 2A) carries a stop-floxed EYFP reporter, which could be construed as a marker of cell-specific VEGF loss. However, we have previously shown that while recombination in stop-floxed reporters can accurately identify the cell population undergoing recombination in a Cre-lox model, they can be poor predictors of recombination of target genes at the single cell level due to the stochastic nature of LoxP recombination in separate genes (Dause and Kirby, 2020).

To be able to reliably differentiate cells that have lost VEGF versus neighboring cells which have not, we used the lentiviral vectors expressing *Vegfa* (or scramble) shRNA plus GFP (Supplemntary Figure 2B,C). We perfused mice 21 days after stereotaxic viral infusion in the adult DG (Figure 8A). The lentiviral vectors showed wide tropism, as expected (Supplementary Figure 4A), and *Vegfa* shRNA reduced VEGF immunoreactivity throughout the DG by 74 ± 12% compared to the scramble control (Supplementary Figure 4B,C). In this model, all cells are exposed to these broad changes in extracellular VEGF, but *Vegfa* shRNA expressing GFP+ cells also lose cell internal VEGF. Neighboring GFP-cells, in contrast, retain that cell autonomous VEGF expression. We found that *Vegfa* shRNA expression led to a significant disruption of vessel association in GFP+ RGL-NSCs and IPCs, with no effect on neighboring GFP-cells (Figure 8C-D). In our previous work (Kirby et al., 2015), we reported that loss of NSPC-VEGF led to RGL-NSC exhaustion at the whole population level, which was characterized by early activation and symmetric exhaustive divisions of RGL-NSCs, causing a surge in IPC production at 21d after knockdown. We therefore quantified the relative percentages of RGL-NSCs and IPCs that were GFP+ and GFP-in shRNA infused mice. We found that *Vegfa* shRNA expression led to significant shift away GFP+ RGL-NSCs in favor of GFP+ IPCs compared to scramble-treated mice, with no such effect in GFP-NSPCs (Figure 8E,F).

**Figure 8:**
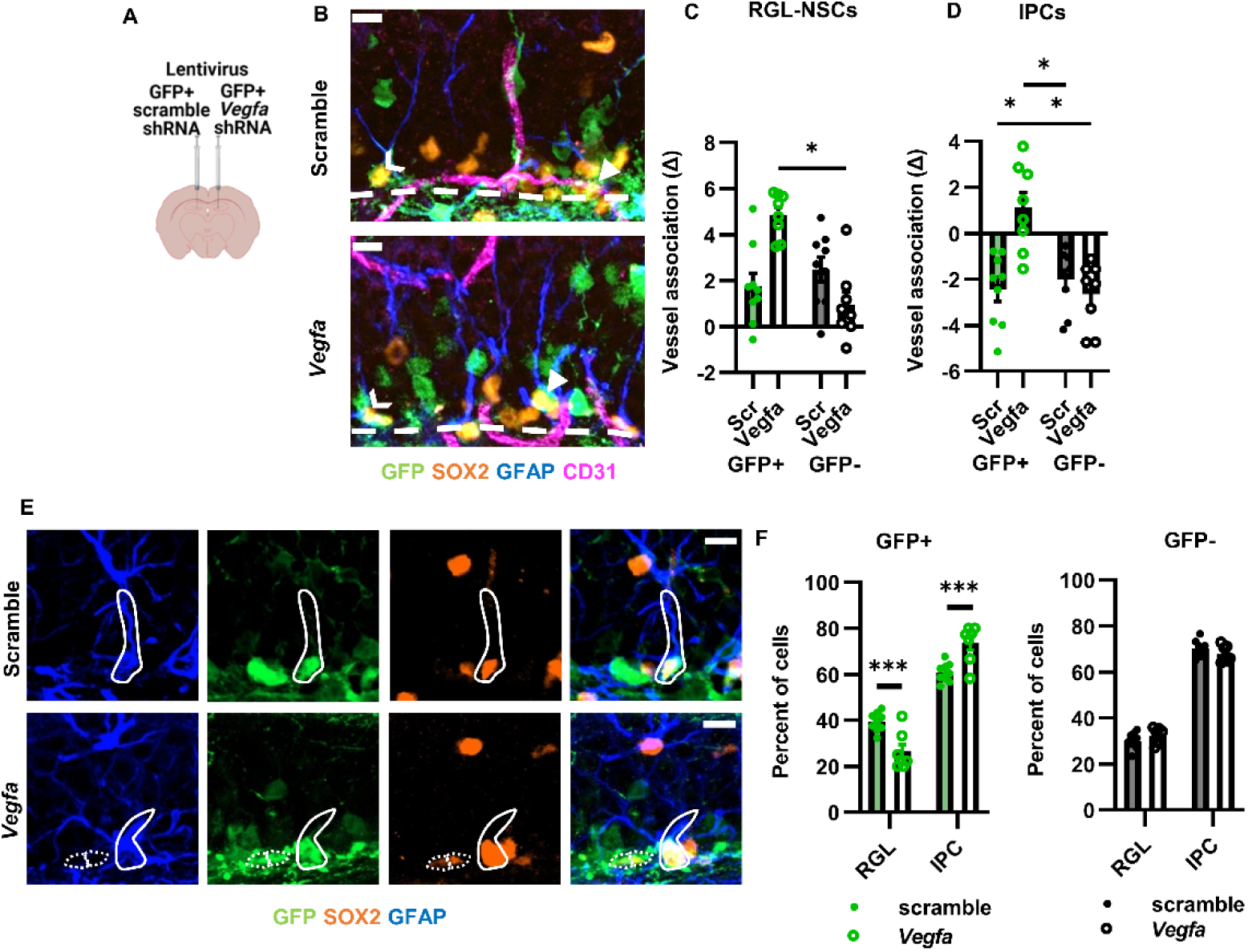
Suppression of VEGF expression cell autonomously reduces vessel association and RGL-NSC maintenance in the adult DG. A) Diagram of experimental design. B) Representative immunofluorescent images of shRNA expressing (GFP+) or non-expressing (GFP-) GFAP+SOX2+ RGL-NSCs and GFAP-SOX2+ IPCs and their association with the CD31+ vasculature 21 days after viral infusion. Chevrons indicate GFP+ RGL-NSCs, arrowheads indicate GFP+ IPCs. C,D) Vessel association measurement of GFP+ or GFP-GFAP+S0X2+ RGL-NSCs (C) or GFAP-SOX2+ IPCs (D) 21 d after viral infusion. E) Representative images of SOX2+GFAP+ RGL-NSCs and IPCs. GFP+ RGL-NSCs have solid outline. GFP+ IPCs have dotted outline. F) Percent of GFP+ and GFP-NSPCs that were GFAP+ SOX2+ RGL-NSCs versus IPCs 21 days after viral infusion. N = 8-9 mice/grp. Mean ± SEM plus individual mice shown. Scale bars = 10 µM. *p < 0.05; *‘*p < 0.001. See also. Supplementary Figure 4.

Our results above suggested that NSC exhaustion may be driven by cell-autonomous VEGF signaling. While the loss of vascular contact could contribute to exhaustion, we also previously reported that NSCs in culture rely on VEGF self-signaling via VEGFR2 to prevent exhaustion (Kirby et al., 2015). To determine whether VEGF directly self-signals in NSCs to maintain self-renewal, we used a neighbor rescue design in cultured NSCs derived from adult male and female DG of VEGF^lox/lox^ mice (Figure 9A). We infected VEGF^lox/lox^ NSCs with a mCherryCre or mCherryonly lentiviral vector at a low multiplicity of infection (MOI), resulting in less than 5% of NSCs showing viral expression (mCherry). High MOI mCherryCre infection led to a significant suppression of VEGF in NSC conditioned media (Figure 9B). At the low MOI, however, extracellular VEGF was not altered (Figure 9C). The low MOI condition therefore provided a model where the infrequent mCherry+ cells have lost cell internal VEGF while still in the presence of abundant extracellular VEGF from neighboring cells. Low MOI mCherryCre treatment led to an acute increase in BrdU+ proliferative cells selectively among mCherry+ cells while mCherryonly expression had no effect on proliferation (Figure 9D,E). There was no effect of mCherryCre versus mCherryonly expression in wildtype (WT) NSCs (Figure 9D,E). When VEGF^lox/lox^ NSCs were grown over multiple passages, a complete and selective loss of mCherryCre+ NSCs was observed while mCherryonly infected NSCs were preserved (Figure 9F,H). There was no effect of mCherryCre versus mCherryonly expression in wildtype (WT) NSCs (Figure 9F-H). These findings suggest that cell autonomous VEGF signaling is necessary to prevent NSC activation and subsequent exhaustion.

**Figure 9:**
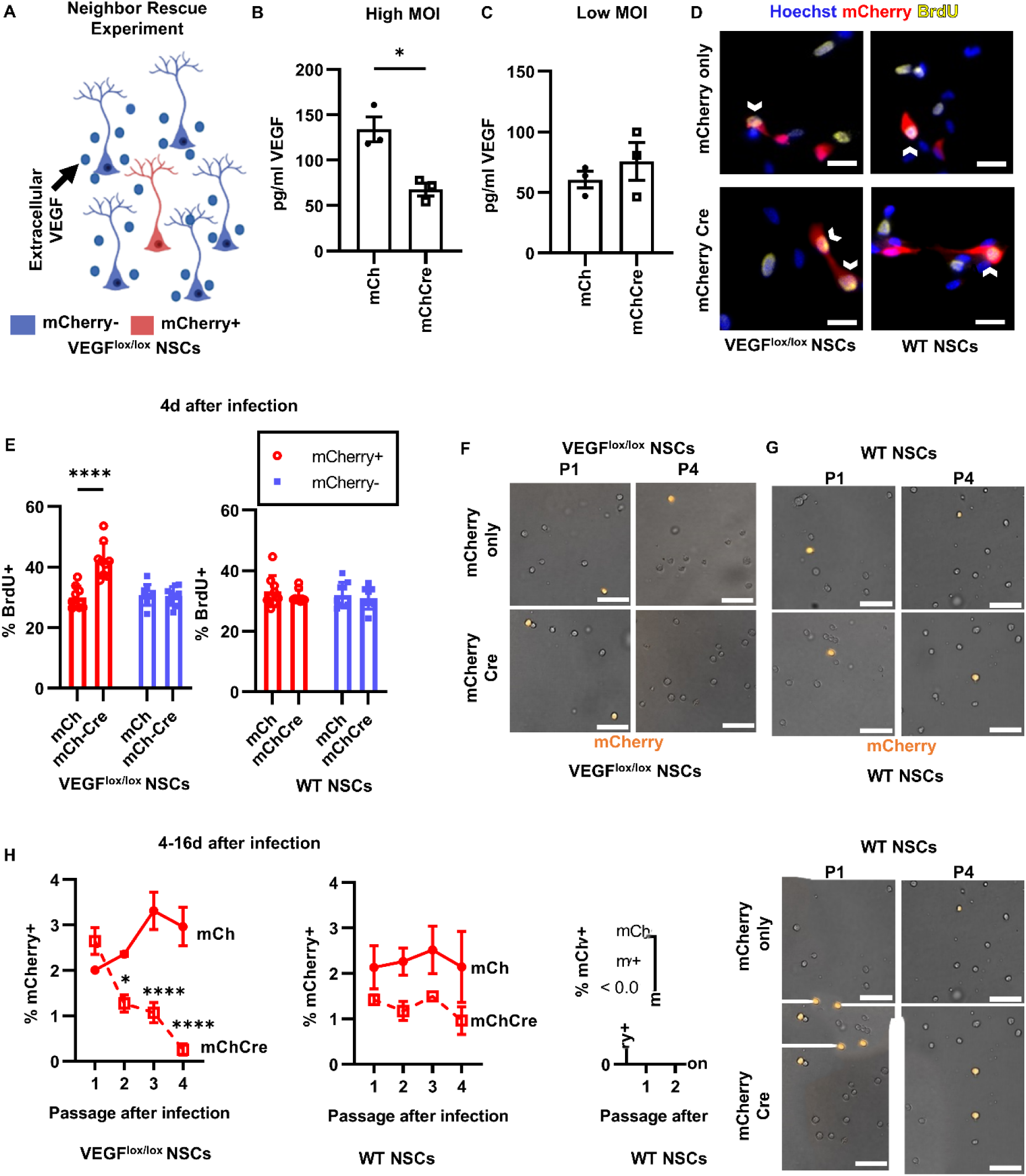
Cell autonomous VEGF signaling maintains NSCs in vitro. A) Diagram of the Neighbor Rescue Experiment. B,C) VEGF concentration (pg/ml) in NSC conditioned media after treatment with high MOI (B) or low MOI (C) mCherry or mCherryCre lentivirus. N = 3 wells/grp; mean ± SEM plus individual wells shown. D) Representative immunofluorescent images of BrdU+ ^cultured^ VEGF^|OX/|OX^ and WT NSCs (Hoechst+) after lentiviral infection (mCherry+). Chevrons indicate BrdU+ mCherry+ Hoechst+ NSCs. E) Percent of mCherry+ or mCherry-VEGF^l0×/l0X^ and WT NSCs that were BrdU+ after low MOI mCherryCre and mCherryonly lentiviral infection. N = 3/grp/exp, 3 exps. Mean ± SEM shown. F) Representative images of cultured VEGF^lox/lθx^ NSCs (brightfield) after lentiviral infection (mCherry+) after 1 and 4 passages. G) Representative images of cultured WT NSCs (brightfield) after lentiviral infection (mCherry+) after 1 and 4 passages. H) Percent of mCherry+ VEGF^lθx/lθx^ and WT NSCs after infection with low MOI mCherryCre or mCherryonly lentiviral vectors. N = 3/grp/exp, 2 exps; mean ± SEM. Scale bars represent D) 10 µM, F,G) 50 µM. *p < 0.05; **p < 0.01; ****p < 0.0001

## Discussion

Cross-talk between adult NSCs and their niche is essential for preservation of neurogenesis through adulthood. The relationship of NSCs and their progenitors with the unique vascular architecture of the adult DG is one of the most prevalent features of this niche. Here, we show that NSPCs are essential to preserving their own proximity to vessels through cell internal activation of VEGF receptors. We also show that cell autonomous VEGF signaling is necessary for preservation of NSC quiescence and prevention of exhaustion independent of vasculature. In vivo, both of these mechanisms could contribute to stem cell maintenance, the former by maintaining blood vessel proximity and the latter via direct promotion of self-renewal (Figure 10). These findings reveal that the maintenance of NSPCs in their vascular niche is not fixed, but rather is actively maintained by NSPCs. They further reveal a surprising mechanism by which NSPCs rely solely on cell internal VEGF receptor activation due to shedding of extracellular facing receptors.

**Figure 10:**
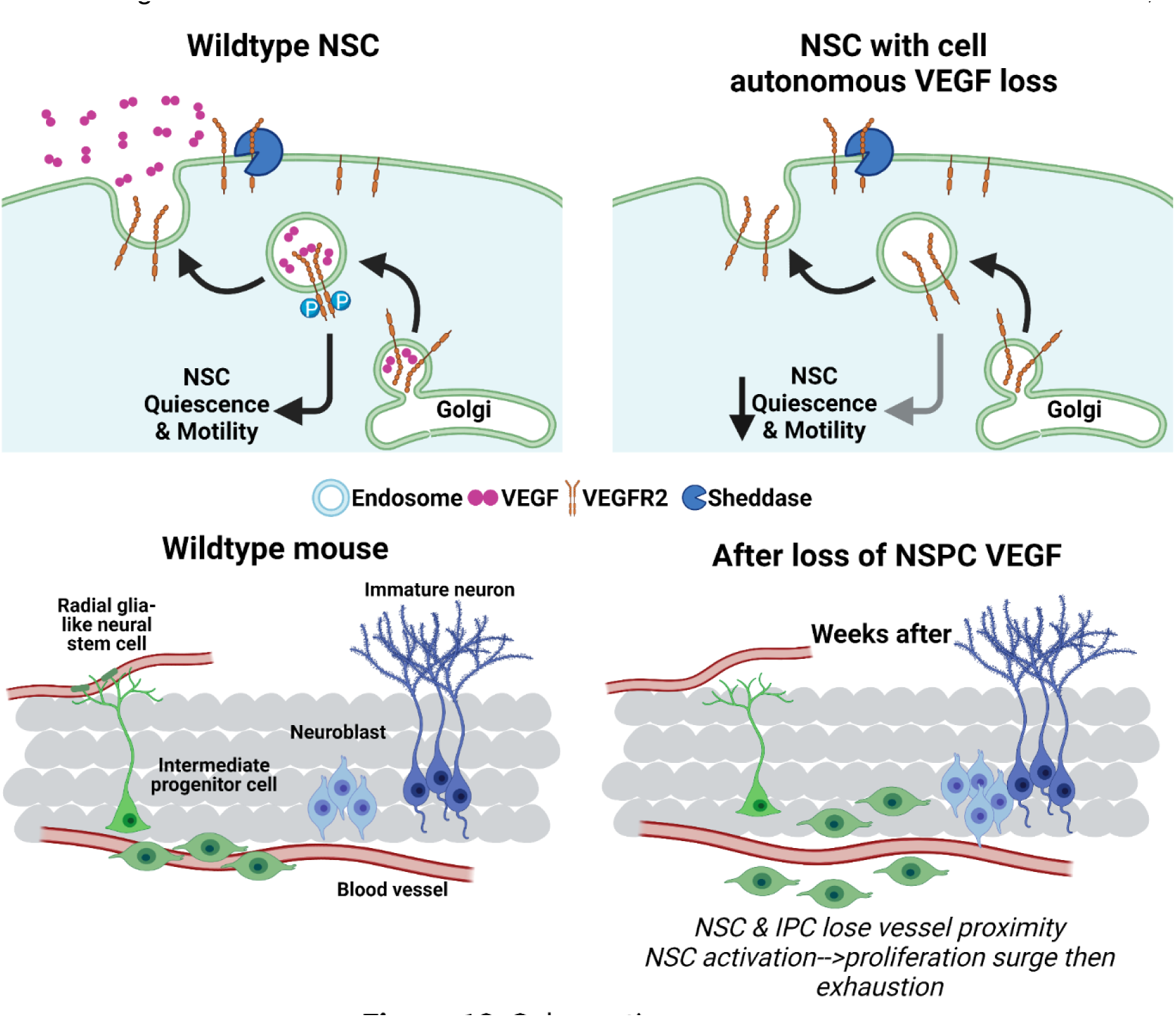
Schematic summary.

The hallmark features of the adult DG neurogenic vascular niche are: 1) density of local blood vessels and 2) NSPC physical apposition to these vessels (Licht et al., 2020a; Moss et al., 2016; Palmer et al., 2000; Sun et al., 2015). There is little information available about the molecular cues that support maintenance of these properties in adulthood. Prenatally, embryonic NSPCs promote de novo vascularization of the CNS via secretion of VEGF, which is a chemoattractant and mitogen for endothelial cells (Gerhardt et al., 2003; Haigh et al., 2003; Raab et al., 2004). VEGF expression in adult brain is typically attributed to mature astrocytes (Bozoyan et al., 2012; Kim et al., 2021b; Licht and Keshet, 2013). However, adult brain vasculature is frequently described as quiescent and VEGF-refractory, obtaining an independence from the reliance on VEGF signaling that characterizes developing vasculature (Wälchli et al., 2023). Our findings align with the concept of adult vasculature being VEGF-refractory. Though we observed a loss of NSPC-vessel association with NSPC-VEGF knockdown, we were unable to detect any changes in vascular architecture. Instead, we observed prominent changes in the motility of adult NSCs, both at the gene expression level in vivo and via direct motility assays in vitro.

An open question from our data is whether IPC motility is impaired by loss of VEGF or if the IPC loss of vessel proximity we observed is a downstream effect of being born from an RGL-NSC that was farther from vessels. IPCs have been shown to be highly motile, using the vasculature as a scaffold to migrate tangentially up to hundreds of microns as they differentiate in to neuroblasts (Sun et al., 2015). IPC motility is typically measured as distance relative to parent RGL-NSC, a measure which technically combines RGL-NSC and IPC movement. However, recent advances with in vivo imaging of RGL-NSCs do not reveal large migratory behaviors in RGL-NSCs, suggesting that the large movements observed are mostly driven by IPCs moving away from parent RGL-NSCs (Bottes et al., 2021; Pilz et al., 2018; Wu et al., 2023). Our scRNAseq data did not reveal many DEGs in IPCs after VEGF loss, suggesting that IPCs may themselves not rely on VEGF signaling to maintain their high level of motility. However, more research is needed to confirm this hypothesis.

The mature DG vascular niche is hypothesized to support adult neurogenesis, in particular via exposure to circulatory factors (Karakatsani et al., 2019). Beneficial blood-borne factors, such as those associated with exercise (Fabel et al., 2003b; Horowitz et al., 2020; Trejo et al., 2001), as well as detrimental factors, such as those associated with old age (Villeda et al., 2011), drive changes in NSPC proliferation that result in altered neurogenesis and correlated improvements or impairments, respectively, in hippocampus-dependent memory function. The unique association of NSPCs with local capillaries is frequently cited as a likely mechanism for how circulating factors drive robust changes in NSPC proliferation and thereby neurogenesis (Kim et al., 2021b; Licht et al., 2020b). Our data show that maintenance of proximity to these factors is not a passive process, but one maintained actively by NSPCs.

In addition to vascular proximity, we also show that cell internal VEGF signaling is necessary to maintain NSCs in vitro, where they are isolated from any endothelial/circulatory factors. VEGF has previously been shown to support survival and/or quiescence through an intracrine loop in several other cell populations, including hematopoietic stem cells, cancer stem cells and endothelia (Dias et al., 2000; Domigan et al., 2015; Gerber et al., 2002). In addition to intracrine signaling loops, however, these cells are typically also sensitive to extracellular VEGF (Wiszniak and Schwarz, 2021). In contrast, we found that DG NSCs are insensitive to extracellular VEGF levels, likely because the extracellular VEGF binding domain of VEGFR2 is cleaved by sheddases on the cell surface. The complete reliance of NSCs on intracrine VEGF appears to be somewhat unique to these cells. The mechanisms by which NSCs in the DG develop reliance on VEGF intracrine signaling to maintain quiescence remain to be investigated, as does the possibility that this shift supports their deepening quiescence over postnatal development (Urbán et al., 2019).

The insensitivity of NSCs to extracellular VEGF that our findings suggest provides substantial resolution for the previously conflicting literature on VEGF and adult neurogenesis. Specifically, past studies suggesting no effect of VEGF loss on adult neurogenesis often relied on methods that neutralized extracellular VEGF (Licht et al., 2011, 2016) while studies suggesting that loss of VEGF signaling suppressed adult neurogenesis often relied on long-term infusion of cell permeable VEGFR2 inhibitors (Fournier et al., 2012; Segi-Nishida et al., 2008; Warner-Schmidt and Duman, 2007). All of these results are consistent with a model where cell internal VEGF is necessary and sufficient to mediate VEGF receptor activation.

There are several limitations to the present study. First, though we have previously shown VEGF protein production by NSPCs as a population (Kirby et al., 2015), our analysis of VEGF and VEGFR2 co-expression in single RGL-NSCs here relies mostly on transcriptional-level analysis. Identification of the cellular source of VEGF protein is complicated by the fact that secreted proteins have short intracellular half-lives and most immunolabeling for VEGF is therefore found in the interstitial space or bound to/inside receptor expressing cells. VEGFR2 antibodies in fixed tissue sections have also proven unreliable (Licht et al., 2016). Indeed, we found that when acutely isolated DG cells were fixed, the VEGFR2 antibody we used lost binding capability even in CD31+ endothelia, which are well-established to express VEGFR2. Second, although we did not detect changes to the vasculature, our subregion analysis may have been too broad to detect subtle changes in vessel location. Additional studies could use live imaging after NSPC-VEGF loss to determine if there is more subtle endothelial migration towards other central sources of VEGF, such as astrocytes. Third, while we saw no changes in occludin expression, these findings do not comprehensively assess changes to BBB permeability. However, VEGF is a permeabilizing factor for the BBB and the BBB in the DG is already reported to be intact (Shen et al., 2004), making it unlikely that loss of VEGF could affect much change under normal circumstances. There is still a possibility that BBB permeability induced by surges of VEGF after an injury (e.g. seizures. stroke) may depend on NSPC-produced VEGF. This possibility remains a topic for future research. Finally, more work is needed to better define the downstream mediators of VEGF signaling in adult DG NSCs. Our single cell sequencing data provides a large dataset that implicates numerous pathways as dependent on NSPC-VEGF signaling. Future research will be necessary to determine the functional roles of those signaling events disrupted downstream of VEGF loss.

In conclusion, determining the key mechanisms by which adult DG NSCs regulate themselves and their microenvironment is an important step in understanding the preservation of the adult hippocampal neurogenic niche. Our findings suggest that VEGF regulates both NSC migration and maintenance, which depends critically on the source of the VEGF ligand. Future studies may examine changes to bidirectional signaling between NSPCs and endothelia to determine if loss of VEGF changes chemokine production in endothelia resulting in a loss of migratory signal for NSCs. They may also investigate how different sources of VEGF or manipulations to VEGFR2 expression may preserve neurogenic capacity, for example to protect against loss of neurogenesis with aging or injury. These findings have important implications when considering how to therapeutically support endogenous adult neurogenesis or exogenous NSC transplants.

## Methods

### Mice

All animal use was in accordance with institutional guidelines approved by The Ohio State University Institutional Animal Care and Use Committee. All mice were 7-10 weeks old at the start of any experimental manipulation. NestinCreER^T2^ mice (Jackson #016261) were crossed with Rosa-stop-floxed-EYFP (Srinivas et al., 2001; Jackson #006148) and VEGF^lox^ mice, provided by Genentech, Inc (Gerber et al., 1999), to create NSPC-specific VEGF knockdown for scRNAseq. Mice were maintained as NestinCreER^T2+/-^;VEGF^lox/wt^;Rosa-EYFP^+/+^ x NestinCreER^T2-/-^; VEGF^lox/wt^;Rosa-LoxSTOPLox-EYFP^+/+^ breeding pairs. NestinCreER^T2+/-^;VEGF^wt/wt^;Rosa-EYFP^+/+^ (WT) and NestinCreER^T2+/-^;VEGF^lox/lox^;Rosa-LoxSTOPLox-EYFP^+/+^ (KD) littermates were used for scRNAseq. Nestin-GFP mice (Mignone et al., 2004; Jackson #033927) were used in flow cytometry. Wild type C57BL/6J male and female mice were purchased from Jackson Laboratory (strain #000664). VEGF-GFP mice were a gift from Brian Seed, Harvard University, Cambridge, MA (Fukumura et al., 1998). All mice were housed in standard ventilated cages, with ad libitum access to food and water throughout all experiments and maintained on a 12h light cycle with lights on at 630h. Male and female mice were represented in approximately equal numbers throughout and no differences in sex were found. We therefore combined them in all presented analysis.

### Tamoxifen Administration

Tamoxifen (TAM; APExBIO) was dissolved in sterile sunflower oil at 20 mg/ml, overnight with agitation at 37°C and stored at +4°C for up to one week in the dark. TAM was injected (180 mg/kg/d, i.p.) for 5 d.

### NSC isolation

NSCs were isolated from adult hippocampus of C57BL/6J or VEGF^lox/lox^ mice (described in Kirby et al., 2015) as described in Babu et al., 2011. Unless otherwise stated, NSCs were maintained on poly-D-lysine (Sigma) and laminin (Invitrogen) coated plates in Neurobasal A media (Invitrogen) with 1x B27 supplement without vitamin A (GIBCO), 1x glutamax (Invitrogen) and 20 ng/ml each of EGF and FGF2 (Peprotech), as per (Babu et al., 2011). No cells were used past passage 20. Two separate lines were used in all experiments, one from 5 pooled C57BL/6J male mice and one from 5 pooled C57BL/6J female mice. No differences between NSCs isolated from males and females were found.

### Endothelial Cells

Human Umbilical Vein Endothelial Cells (HUVEC, Invitrogen) were maintained as per manufacturer instructions on uncoated tissue culture plates. bEnd.3 cells (ATCC) were maintained in Dulbecco’s Modified Eagle’s Medium (ATCC 30-2002) with 10% fetal bovine serum (Fisher) at 37 °C, 5% CO_2_ in tissue culture dishes.

#### RNAscope in situ hybridization and immunohistochemistry

WT C57BL/6J mice were transcardially perfused with ice cold PBS followed by 4% PFA. Brains were harvested and fixed overnight at 4°C in 4% PFA before serial overnight equilibration in 10%, 20%, and 30% sucrose. Fixed tissue was snap frozen in OCT in a dry ice/100% ethanol bath and stored at −70°C. 12 µm cryosections, 1 section per slide, were prepared with a cryostat and stored at −70°C with desiccant until staining. RNA in situ hybridization was performed according to manufacturer recommendations for using fixed frozen tissue samples in the RNAscope Multiplex Fluorescent v2 Assay (Advanced Cell Diagnostics) with the following modifications to enable concurrent immunohistochemical staining. The pretreatment steps were replaced with a 15 min modified citrate buffer (Dako) antigen retrieval step in a steamer at 95°C. Additionally, the protease III step was excluded to enable subsequent immunohistochemical staining. Probes for mouse *Vegfa* (Mm-Vegfa-ver2; ACD) and mouse *Kdr* (Mm-Kdr-C2; ACD) RNA were hybridized to tissue before subsequent immunohistochemical staining for GFAP protein. Immunostaining for GFAP was conducted as described in (Immunohistochemical tissue processing) with the following exceptions. Blocking was performed with 10% normal donkey serum in 0.1M tris buffered saline (TBS)-1% bovine serum albumin (BSA). Antibody incubations were performed in TBS-1% BSA. All washes were performed with TBST. DAPI provided by the RNAscope Multiplex Fluorescent kit was used for nuclear counterstaining. All images were acquired with the Zeiss Axio Observer Z1 microscope with Apotome for optical sectioning using a 20x air objective. Full z-stacks were acquired for analysis.

### RNAscope analysis

RGL-NSCs and astrocytes were identified based on the morphology of GFAP+ apical or stellate processes, respectively, extending from a Hoechst+ nucleus in 1 µm z-stack images from n = 4 mice. *Vegfa* puncta were found almost exclusively in the nucleus in both cell types and were counted manually throughout the depth of the nucleus. *Kdr* puncta were also counted within RGL-NSC Hoechst+ nuclei.

### Dentate gyrus isolation and flow cytometry

DG isolation and flow cytometry was used to identify VEGFR2 surface expression. For VEGFR2 surface expression, 8-10 week old Nestin-GFP adult mice (n = 5) were anesthetized and perfused with HBSS without Ca^2+^/Mg^2+^. Following perfusions, brains were removed and put in ice cold HBSS until dissection. Brains were then bisected along the midsagittal line and overlaying diencephalic structures were removed. DGs were dissected under a dissection microscope (Zeiss) according to Hagihara et al., 2009. In brief, the DG was separate from the surrounding tissue with a beveled syringe needle and placed in ice cold HBSS without Ca2+/Mg2+. DGs were then mechanically dissociated with sterile scalpel blades then repeatedly with mortar and pestle in douncing buffer on ice. Dissociated cells were collected by centrifugation at 500 *g* for 5 min before resuspending in HBSS without calcium/magnesium. Cells were then filtered through a 35 μm nylon filter before staining with fluorescent antibodies (Reagents and Resources Table) on ice for 30 min. During the last 10 min of staining, Hoechst dye was added for live/dead discrimination. All cells were washed twice following staining and immediately processed on the FACS Aria III (BD Biosciences, Franklin Lakes, NJ, United States). The data were analyzed using FlowJo^TM^ v10.8.1 software (BD Life Sciences) and NSCs or endothelial populations were identified based on fluorescent markers with the GFP+, Glast+, CD31-, CD45-,CD11b-, O1-, and O4-live cells were designated as NSCs while CD31+, CD45-, CD11b-, O1-, and O4-live cells were designated as endothelia. NSCs and endothelia were then analyzed for VEGFR2 expression.

### Fluorescence Activated Cell Sorting (FACS) for 10x Genomics Single Cell Sequencing and Analysis

NestinCreER^T2+/-^; VEGF^lox/lox^; ROSA-LoxSTOPLox-EYFP^+/+^ (KD; n = 5) and NestinCreER^T2+/-^; VEGF^wt/wt^; ROSA-LoxSTOPLox-EYFP^+/+^ (WT; n = 5) littermates were submitted to tamoxifen injections (described above) and 3 weeks later their DGs were dissected, as described above. DGs were then mechanically dissociated with sterile scalpel blades then repeatedly with mortar and pestle in douncing buffer on ice. Dissociated cells were collected by centrifugation at 500 *g* for 5 min. Cells were then in an HBSS without Ca^2+^/Mg^2+^/Percol solution to form a gradient and spun at 450 *g* for 15 min at room temp. The supernatant was removed, and the remaining cells were resuspended in a solution containing HBSS with 0.04% BSA. Cells were then filtered through a 35 μm nylon filter before staining with Hoechst dye on ice for 10 min for live/dead discrimination. All cells were washed twice following staining and immediately sorted on the FACS Aria III (BD Biosciences, Franklin Lakes, NJ, United States) where EYFP+ cells were collected for scRNAseq in the HBSS 0.04% BSA buffer. Of those, ∼10,000 cells per genotype were loaded onto the 10× Genomics single cell sequencing platform using the standard kit. The 3′ RNA-seq library was sequenced using paired-end 150 bp approach on an Illumina NovaSeq 6000 sequencer. CellRanger 7.0.1 (Zheng et al., 2017) was used to demultiplex, align, and deduplicate sequencing reads in BCL files. Single-cell data in feature-barcode matrices were then processed using Seurat v4.1.1’s default pipeline (Hao et al., 2021) to identify unsupervised cell clusters and generate a uniform manifold approximation and projection (UMAP) plot. In brief, cells were filtered to exclude multiplets and damaged cells by excluding cells with unique feature count <2,000 or >5,000 and a mitochondrial gene expression of >2.0%. From ∼10,000 cells loaded per genotype, ∼5,500 were recovered in both genotypes, yielding a net capture rate of 55%. Data were then log normalized with default scale factor of 10,000 and integrated into one file containing both KD and WT samples. The data underwent linear transformation (ScaleData function) and PCA was run on the scaled data, followed by FindNeighbors and FindClusters. UMAPs were then created using the RunUMAP function. DEGs defining clusters (regardless of treatment) and defining treatments (regardless of cluster) were generated using the FindAllMarkers function, which uses a default of Wilcoxon rank sum test, unless otherwise noted. Adjusted *P*-values are Bonferroni-corrected using all features in the dataset.

### In vitro treatment

SU5416 and SU1498 (Sigma) were dissolved in DMSO (10 mM) and stored at −20°C until use. SU5416 was used at 25 µM unless otherwise noted and SU1498 was used at 2.5 µM. VEGF nAb in sterile PBS (B20-4.1.1) was provided by Genentech Inc. (Pan et al., 2007) and used at 10 µg/ml unless otherwise noted. Mouse recombinant VEGF164 (R&D Systems) was dissolved in sterile PBS and stored at −20°C. It was used at 10 ng/ml unless otherwise noted.

### Immunoblotting

For p/AKT and p/Erk immunoblotting NSCs were plated at a density of 300,000 cells/well in 6-well plates and treated at ∼70% confluency. HUVECs were plated on uncoated 12 well plates at a density of 50,000 cells/well and maintained for 2d then serum starved for 6 h before treatment. NSC and HUVEC monolayers were lysed with RIPA buffer (Pierce) with 1x Halt Protease and Phosphatase Inhibitor (Thermo Scientific) on ice for 10 min. Scraped lysates were freeze-thawed 3 times and then centrifuged at 14000 rpm for 10 min at 4°C. Total protein concentration of the supernatant was quantified using a BCA kit (Pierce). Lysates were run on 4-12% bis-tris gels in 1x NuPage MOPS buffer (Invitrogen) at 120 V for 2 h. Gels were transferred to nitrocellulose membrane overnight in 20% methanol in NuPage Transfer Buffer (Invitrogen) at 4°C. Membranes were blocked in 5% milk in 0.1M TBS with 0.5% Tween20 (TBS-t) then incubated in primary antibody overnight at 4°C in 5% milk in TBS-t. Secondary was applied for 1 h at room temperature in 5% milk in TBS-t. Proteins were visualized and quantified on a LICOR Odyssey Clx infrared imaging system. Quantification was taken as the ratio of phosphorylated protein to its unphosphorylated form.

For pVEGFR2 detection, NSCs were plated in a 10cm dish format and grown to confluency before treatment with SU5416 or control for 10 min before harvesting for lysates as above. Due to low expression/detection of pVEGFR2 in NSC samples, VEGFR2 was first immunoprecipitated (i.p.) from NSC lysates using the Dynabeads^TM^ Immunoprecipitation Kit (Thermo Scientific), following all manufacturer instructions. VEGFR2 immunoprecipitants were then run on 4-12% bis-tris gels and transferred to nitrocellulose membranes as above probed for pVEGFR2. Membranes were blocked in 5% BSA in TBS-t then incubated in primary antibody overnight at 4°C in 5% BSA in TBS-t. Secondary was applied for 1 h at room temperature in 5% BSA in TBS-t. pVEGFR2 expression was visualized and quantified on a LICOR Odyssey Clx infrared imaging system.

### Lentiviral vector production

mCherry-Cre (Addgene no. 27546) and mCherry-only vectors are described in Kirby et al., 2015. Plasmids were packaged in vesicular stomatitis virus-glycoprotein G (VSV-G) lentivirus by the Cincinnati Children’s Hospital Viral Vector Core. VEGF and scramble shRNA lentiviral vectors are described in (Mosher et al., 2012) and were packaged in VSV-G lentivirus by either Vigene Biosciences or the Stanford Gene Vector and Virus Core.

### Neighbor rescue experiment

Adult DG NSCs were grown in standard media on uncoated, tissue culture plates, allowing them to form spheres. On day 1, 5000 NSCs derived from DG of 6-week-old VEGF^lox/lox^ mice or VEGF^wt/wt^ mice were infected with mCherry-Cre or mCherry-only lentiviral vectors titered to result in <5% total mCherry+ cells. 4 days after infection, short term neighbor rescue experiment NSCs were incubated with 20 µM 5-bromo-2’-deoxyuridine (BrdU) (Sigma) for 2 hours then fixed with 4% paraformaldehyde prior to immunohistochemical processing. Long term neighbor rescue NSC spheres were dissociated every 4d and freshly plated single cells were imaged live to quantify the percent of mCherry+ cells among those identified by brightfield contrast. Images were quantified by a blinded observer.

### VEGF ELISA of in vitro VEGF^lox/lox^ conditioned media

Adult DG VEGF^lox/lox^ NSCs were infected with mCherry-Cre or mCherry-only lentiviral vectors titered to result in <5% mCherry+ cells (low) or >50% mCherry+ cells (high). After 4d, conditioned media was collected and centrifuged 1000g for 5 min. Conditioned media supernatant was extracted and processed with the Mouse VEGF DuoSet ELISA (R&D Systems, DY493-05) according to manufacturer instructions.

### shRNA Lentiviral Infection in vitro

Adult Wt C57BL6/J mouse DG derived NSCs were plated on coated 96-well plates at a density of 10,000 cells/well then infected with lentiviral vectors containing a VEGF shRNA or a scramble shRNA (described in Mosher et al., 2012) at a MOI of 50. Media was changed after 24h. After 3 d, NSPCs were transferred to sphere conditions on uncoated plates at 5000 cells/well and fed FGF2/EGF every 2d. Sphere number and size was quantified using a CellAvista automated microscope system (Roche). Conditioned media was collected from supernatant at first passage and assayed with the Mouse VEGF DuoSet ELISA (R&D Systems) according to manufacturer instructions.

## Scratch Assay

For SU5416 treatment, NSCs were plated into a 24-well plate coated with laminin and PDL at a density of 50,000 cells per well. SU5416 was dissolved in DMSO (10 mM) and stored at −20°C until use. SU5416 was used at 25 µM. After 24 hours, a scratch was performed with a sterile pipet tip. The scratch was then imaged at 1h, 4h, and 8h to observe NSC migration, while incubating at 37 °C between the time points. For lentiviral infection, NSCs were plated into a 24-well plate coated with laminin and PDL at a density of 20,000 cells per well. 24 hr later cells were treated with *Vegfa* or scramble virus. After 72 hours, a scratch was performed as above and images 1h, 4h and 8h afterwards to track NSC migration.

### Stereotaxic Surgery

Mice were anesthetized by inhalation of isoflurane (Akorn, 5% induction, 1-2% maintenance) in oxygen and mounted in the stereotaxic apparatus (Stoelting). Ocular lubricant (Puralube) was placed over the eyes to prevent evaporative dry eye. Following sterilization with alcohol (Fisher) and betadine swabs (Fisher), the skull was exposed and the lambda and bregma sutures were aligned in the same horizontal plane. A small bur hole was drilled in the skull and an automated injector (Stoelting) with a Hamilton syringe (Hamilton) was lowered to the injection depth at a rate of −1.0 mm/min. Mice were injected with 0.5 μL of scramble control shRNA virus into one hemisphere and 0.5 μL of *Vegfa* shRNA virus into the contralateral hemisphere at a rate of 0.1 μL/min. The injection coordinates from bregma were: anterior/posterior -1.9 mm, medial/lateral ±1.6 mm, -1.9 mm dorsal/ventral from dura. Post-surgery, the incision was sealed with tissue adhesive (3M) and the mouse was given saline (Hospira) and carprofen (Zoetis) injection i.p. After 21d, mice were perfused for immunohistochemical processing.

### EdU and BrdU labeling

21d after TAM administration mice were injected with 5-bromo-2’-deoxyuridine (BrdU, 150 mg/kg, IP) (Sigma) and sacrificed 2 hours later for tissue processing. In virus-treated mice, after 21d, mice were injected with 5-Ethynyl-2’-deoxyuridine (EdU) (Click Chemistry Tools) dissolved in physiological saline (Hospira) (150 mg/kg, IP) and sacrificed 2 hours later for tissue processing.

### Immunohistochemical processing

#### Cultured NSCs

Adherent NSCs were fixed with 4% paraformaldehyde then rinsed with PBS 3x before incubating in a blocking solution containing 1% normal donkey serum (Jackson ImmunoResearch) and 0.3% Triton X-100 (Acros) in PBS. Cells were then incubated in primary antibody diluted in blocking solution overnight at 4 °C. The following day, after 3 rinses in PBS, cells were incubated in secondary antibodies diluted in blocking solution for 2 hours. If a biotinylated secondary was used, a fluorophore-conjugated tertiary was applied for 1h diluted in PBS before rinsing and nuclei counterstaining with Hoechst (10 min, 1:2000 in PBS) (Invitrogen). If proceeding for BrdU labeling, the cells were rinsed and fixed with 4% paraformaldehyde in 0.1M PB for 10 min, rinsed with PBS 3x and incubated with 2N HCl for 30 min at 37 °C. After 3 PBS rinses and 30 min incubation in blocking, cells were incubated in BrdU primary antibody diluted in blocking solution overnight at 4 °C. The next day, cells were rinsed 3x with PBS and exposed to a secondary antibody diluted in blocking solution for 2h before Hoechst counterlabeling. Cells in 96-well plates were imaged immersed in PBS using 10x magnification while those on chamber slides were coverslipped with Prolong Gold Antifade Mountant (Fisher) and imaged using 40x oil magnification, both with a Zeiss apotome digital imaging system (Zeiss).

#### Brain Sections

Brains for immunolabeling were harvested following perfusion with ice-cold PBS followed by fixation in 4% paraformaldehyde overnight at 4 °C. After equilibration in 30% sucrose in PBS, 40 μm coronal brain sections were obtained in 1 in 12 series on a freezing microtome (Leica) and stored in cryoprotectant at −20 °C. Sections were rinsed 3x in PBS and incubated in a blocking solution containing 1% normal donkey serum and 0.3% Triton X-100 (Acros) in PBS before incubation in primary antibodies. The next day, cells were rinsed 3x with PBS and exposed to a secondary antibody diluted in blocking solution for 2h. EdU labeling was performed according to manufacturer recommendations for using fixed frozen tissue samples to enable concurrent immunohistochemical staining (Click Chemistry Tools) before the immunolabeling protocol. BrdU immunolabeling was performed after other immunolabeling was complete. Labeled sections were fixed in 4% paraformaldehyde for 10 min at room temperature, rinsed and then incubated with 2N HCl for 30 min at 37 °C. Sections were then rinsed, blocked and incubated in anti-BrdU primary and appropriate secondaries as above. The DG of the hippocampus was imaged in 1 μm Z-stacks at 20x magnification using a Zeiss apotome digital imaging system (Zeiss).

### Immunofluorescent image quantification

In TAM treated mice applications, RGLs were identified by GFAP+/EYFP+ colocalization and GFAP+ radial processes extending from the SGZ towards the inner molecular layer, while MCM2+/EYFP+ cells in the SGZ layer were identified as IPCs. BrdU+ nuclei were counted in the subgranular zone and granular cell layer. Neuroblasts were identified as DCX+ cells in the SGZ with a bipolar morphology. Immature neurons were identified as DCX+ cells with a primary dendrite extending through the granular cell layer. Endothelia were identified by CD31+.

Distance to vasculature was measured as the distance from the middle of a cell body to the nearest CD31+ vessel. Random distances for vessel associations were measured by sampling the distance of random Hoechst+ cells in the middle of the SGZ to the vasculature. Vessel association was calculated by the mean distance of a cell to the vasculature minus the mean distance of a “random cell”. The SGZ was defined as the zone spanning 2 cell body widths between the dense granular cell layer and the hilus. CD31+ cell density and Occludin/CD31 overlap were performed calculated by thresholded are in ImageJ by a blind observer.

In other applications, NSCs were identified by GFAP+/SOX2+ colocalization and GFAP+ radial processes extending from the SGZ towards the inner molecular layer, while GFAP-/SOX2+ cells in the SGZ layer were identified as IPCs. In mice infused with shRNA lentivirual vectors, NSPCs were identified as above and then categorized as GFP+ or GFP-within the SGZ, yielding a percent of GFP+ or GFP-NSPCs that were RGL-NSC or IPC phenotype. For RNAscope alone, GFAP+ apical process alone was used (without SOX2) due to limitations in the number of separate fluorescent wavelengths available. The SGZ was defined as the zone spanning 2 cell body widths between the dense granular cell layer and the hilus. Cell counts were performed manually in Zen by a blind observer.

### CRISPRi VEGFR2 knockdown for immunolabeling

bEnd.3 cells or NSCs were plated on uncoated (bEnd.3) or on PDL/laminin-coated (NSCs) glass 8-chamber slides (Fisher) and allowed to adhere for 24h. After 24h, cells were transfected with CRISPRi expressing plasmids using Lipofectamine 2000 (Fisher), according to manufacturer instructions. Media was changed to remove lipofectamine/DNA 5h later and replaced with standard growth media for each respective cell type. Two days later, cells were fixed with 4% paraformaldehyde.

### Plasmid construction and sgRNA design

The CRISPR interference (CRISPRi) lentivirus construct was modified from pLV hU6-sgRNA hUbC-dCas9-KRAB-T2a-GFP (Addgene #71237, a gift from Charles Gersbach, Thakore et al., 2015), which expresses all necessary CRISPRi machinery (both the dCas9-KRAB and sgRNA) from the same plasmid. The UbC promoter was replaced with a Nestin regulatory element comprised of the Nestin promoter and second intronic enhancer from (Keyoung et al., 2001) to create pLV hU6-sgRNA Nestin-dCas9-KRAB-T2a-GFP. Additionally, two Esp3I recognition sites were placed after the U6 promoter for restriction cloning of sgRNA insert sequences. The insert sequences were synthesized as single strand oligonucleotides (Integrated DNA Technologies) in the form of 5’ GGACG(N)_20_ 3’ and 5’ AAAC(N’)_20_C 3’ where (N)_20_ refers to the sequence of the sgRNA and (N’)_20_ is the reverse complement. These oligonucleotides were annealed together and ligated with Esp3I-digested pLV hU6-sgRNA Nestin-dCas9-KRAB-T2a-GFP to obtain the final constructs for lentivirus packaging. The sgRNAs targeting the region −50 to +300 bp relative to the transcriptional start sequence of *Kdr* were designed using the CRISPRi function of the Broad Institute GPP portal (https://portals.broadinstitute.org/gpp/public/analysis-tools/sgrna-design). The top 5 ranked sgRNAs returned by the GPP tool were chosen and a non-targeting sgRNA (Chiou et al., 2015) was used as control.

### VEGFR2 cell surface and intracellular immunolabeling and quantification

To label all VEGFR2, blocking solution was 1% normal donkey serum, 0.3% tritonX 100 in PBS. To label only cell surface VEGFR2, the triton 100X was omitted but all other procedures were identical. 3 random view fields were captured for VEGFR2 intensity quantification using thresholded area and intensity within cell bodies using Image J.

### TAPI1 treatment

NSCs derived from 6-week-old Wt C57BL6/J DG were plated at 20,000 cells/well on PDL/laminin-coated glass 8-chamber slides (Fisher 08-774-26) and allowed to adhere for 24h. TAPI1 (Fisher 61-621) was added to media at 0-80 µg/ml. NSCs were fixed 24h later. For TAPI rescue experiments DG NSCs were treated with 160 µg/ml 24 hours before 10 min of VEGF164 treatment (10 ng/ml, R&D Systems).

### CRISPRi VEGFR2 knockdown for qPCR

bEnd.3 cells were plated on 12-well tissue culture plates (VWR #82050-930) and allowed to grow to 80% confluency before transfection with CRISPRi plasmids using Lipofectamine 2000 (Fisher), according to manufacturer instructions. 18h later, cells were harvested by scraping in 0.25% Tryspin-EDTA with phenol red (Fisher), then centrifuged 1000 g for 5 min. RNA was isolated from cell pellets using the Aurum Total RNA Mini kit (Bio-Rad). Total RNA was quantified using a BioTek Epoch Microplate Spectrophotometer and 250 ng RNA was converted to cDNA using the iScript cDNA synthesis kit (Bio-Rad) and a ThermoFisher Applied Biosystems 2720 Thermal Cycler. cDNA was amplified with gene-specific primers and SsoAdvanced Universal SYBR Green Supermix in a Bio Rad CFX96 Touch Real-Time PCR Detection System. Melt curves were used to confirm purity of the amplified product and Ct value was normalized to houskeeping gene *Hprt*. ΔΔCt values were used to obtain fold change over non-target control average.

Hprt: PrimerBank ID: 96975137c1

Forward primer: agtcccagcgtcgtgattag

Reverse primer: tttccaaatcctcggcataatga

Kdr: From Kirby et al., 2015

Forward primer: atctttggtggaagccacag

Reverse primer: ccatgatggtgagttcatcg

### Diagram creation

All diagrams in this manuscript were created with <u>BioRender.com.</u>

### Quantification and Statistical Analysis

Statistics were performed as described in each figure legend. Statistical details for each data panel except for scRNAseq is in Table S2. In general, if pairs of groups were compared, t-tests were used. If more than 2 groups were compared with 1 factor, one way ANOVA was used followed by error-corrected posthoc tests. If groups with 2 factors were compared, two way ANOVA was used followed by error-corrected posthoc tests. Three-way ANOVAs were used to compare virus x day x GFP expression interaction in viral infected tissue sections. After confirmation of significant 3-way interaction, two-way ANOVAs were used within GFP+ or GFP-cell populations to confirm a significant virus x day interaction followed by Sidak’s multiple comparisons post hoc tests. All analyses were performed using Prism (v9.0; GraphPad Software) and p<0.05 was considered significant. Single cell RNAseq analysis is described in the Single Cell section above.

## Reagents and Resources Table

| REAGENT or RESOURCE | SOURCE | IDENTIFIER |
| --- | --- | --- |
| **Antibodies** |  |  |
| Cell Signaling Technology, 2920s, Akt (pan) (WB; 1:100) | Cell Signaling Technology | Cat#2920<br>RRID:<br>AB_1147620 |
| Cell Signaling Technology phospho-Akt (ser473) (WB; 1:1000) | Cell Signaling Technology | Cat#4060<br>RRID:<br>AB_2315049 |
| VEGF Receptor 2 (D5B1) Rabbit mAb<br>(i.p.; 1:500) (IF; 1:50) | Cell Signaling<br>Technology | Cat#9698<br>RRID: |
| IRDye® 680RD Goat anti-Mouse IgG (H + L), 0.1<br>mg [P/N 925-68070] (WB; 1:10,000) | VWR | Cat#: 102971-020<br>RRID: |
| IRDye® 800CW Goat anti-Rabbit IgG (H + L), 0.1<br>mg [P/N 925-32211] (WB; 1:10,000) | VWR | Cat# 103011-494<br>RRID: |
| Hoechst 33342<br>(IF; 1:2000) (Flow; 1:5000) | Fisher | Cat#H3570<br>RRID: |
| Anti-Glial Fibrillary Acidic Protein, Clone: GA5<br>(IF; 1:1000) | Sigma | Cat#MAB360<br>RRID:<br>AB_11212597 |
| Anti-mCherry antibody<br>(IF; 1:500) | Abcam | Cat#ab167453<br>RRID:<br>AB_2571870 |
| Anti-GFP Antibody<br>(IF; 1:1000) | abcam | Cat#ab6673<br>RRID: AB_305643 |
| BrdU Antibody<br>(IF; 1:500) | Bio-Rad | Cat# MCA6114<br>RRID: |
| Goat anti-VEGF164<br>(Neutralization; 10µg/ml) (IF; 1:50) | R&D systems | Cat#AF-493<br>RRID: AB_354506 |
| SOX2 Rat anti-Human, Mouse, Clone: Btjce<br>(IF; 1:1000) | eBioscience | Cat# 50-112-9095<br>RRID: |
| Anti-NeuN, clone A60<br>(IF; 1:500) | Millipore | Cat#MAB377<br>RRID: |
| Rat anti-VEGFR2<br>(IF; 1:40) | R&D systems | Cat#MAB4432<br>RRID:<br>AB_2298772 |
| Donkey anti-Goat IgG (H+L) Cross-Adsorbed<br>Secondary Antibody, Alexa Fluor 488 (IF; 1:500) | Fisher | Cat#A-11055<br>RRID:<br>AB_2534102 |
| Donkey anti-Rabbit IgG (H+L) Highly Cross-Adsorbed Secondary Antibody, Alexa Fluor 555<br>(IF; 1:500) | Fisher | Cat#A-31572<br>RRID: AB_162543 |
| Donkey anti-Goat IgG (H+L) Cross-Adsorbed<br>Secondary Antibody, Alexa Fluor 555 (IF; 1:500) | Fisher | Cat#A-21432<br>RRID:<br>AB_2535853 |
| Donkey anti-Mouse IgG (H+L) Highly Cross-Adsorbed Secondary Antibody, Alexa Fluor 647<br>(IF; 1:500) | Fisher | Cat#A-21447<br>RRID: AB_141844 |
| Donkey anti-Rat IgG (H+L) Highly Cross-Adsorbed<br>Secondary Antibody, Alexa Fluor 594 (IF; 1:500) | Fisher | Cat#A-21209<br>RRID:<br>AB_2535795 |
| IgG (H+L) Highly Cross-Adsorbed Donkey anti-Rabbit, Alexa Fluor™ 350 (IF; 1:500) | Fisher | Cat#A-10039<br>RRID:<br>AB_2534015 |
| PE/Cyanine7 anti-mouse CD31 Antibody<br>(Flow; 1:25) | BioLegend | Cat#102418<br>RRID: AB_830757 |
| GLAST (ACSA-1) Antibody, anti-human/mouse/rat<br>(Flow; 1:100) | Miltenyi Biotec | Cat#130-118-483<br>RRID:<br>AB_2733472 |
| Recombinant APC Anti-VEGF Receptor 2 antibody (Flow; 1:100) | abcam | Cat# ab275611<br>RRID: |
| Oligodendrocyte Marker O1 Alexa Fluor® 700-conjugated Antibody (Flow; 1:100) | R&D systems | Cat# FAB1327N<br>RRID: |
| Oligodendrocyte Marker O4 Alexa Fluor® 700-conjugated Antibody (Flow; 1:100) | R&D systems | Cat# FAB1326N<br>RRID: |
| Alexa Fluor® 700 anti-mouse CD45 Antibody (Flow; 1:100) | BioLegend | Cat# 103127<br>RRID: |
| Alexa Fluor® 700 anti-mouse/human CD11b Antibody (Flow; 1:100) | BioLegend | Cat# 136405<br>RRID: |
| **Chemicals, peptides, and recombinant proteins** |  |  |
| SU5416 (25 µM) | Sigma | Cat# S8442 |
| SU1498 (2.5 µM) | Sigma | Cat# SML1193 |
| VEGF nAb (10 µg/mL) | Genentech Inc | Pan et al., 2007 |
| Mouse recombinant VEGF164 (10 ng/ml) | R&D Systems | Cat# AF493SP |
| Animal-Free Recombinant Human EGF, PeproTech (AF10015-500UG) | VWR | Cat#: 10781-694 |
| PeproTech Recombinant Human FGF-Basic (154A.A) (100-18B-250UG) | VWR | Cat#: 10771-938 |
| Lipofectamine 2000 | Invitrogen | Cat#: 11668-019 |
| Gibco™ HBSS without Calcium, Magnesium or Phenol Red | Fisher | Cat#: 14-175-095 |
| Gibco™ (Phosphate Buffered Saline) Solution, pH 7.4, 1X | Fisher | Cat#: 10-010-049 |
| Albumin, Bovine Serum, Fraction V, Fatty Acid-Free, Nuclease- and Protease-Free | Millipore Sigma | Cat#: 126609 |
| TRITON™ X-100 | Thermo Scientific Chemicals | Cat#: 21568 |
| 5-bromo-2'-deoxyuridine | Sigma | Cat# B5002 |
| 5-Ethynyl-2'-deoxyuridine | Click Chemistry Tool Kit | Cat#1324 |
| TAPI1 | Fisher | Cat#61-621 |
| **Critical commercial assays** |  |  |
| RNAscope Multiplex Fluorescent Assay v2 | ACD Biotechnne | Cat#: 323136 |
| RNAscope probe Mm-Vegfa-ver2 | ACD Biotechnne | Cat#: 412261 |
| RNAscope probe Mm-Kdr-C2 | ACD Biotechnne | Cat#: 414811-C2 |
| Mouse VEGF DuoSet ELISA | R&D Systems | Cat# DY493-05 |
| Click-&-Go EdU 647 Cell Proliferation Assay Kit | Click Chemistry Tool Kit | Cat# 1329 |
| **Deposited data** |  |  |
| Single Cell RNAseq Data Set |  | GEO#: GSE220871 |
| **Experimental models: Cell lines** |  |  |
| Cultured Neural Stem Cells | Babu et al., 2011 | doi:<br>10.3389/fnins.2011.00089 |
| bEnd.3s | ATCC | Cat#2299<br>RRID: |
| HUVECs | Invitrogen | Cat# C0035C<br>RRID: |
| **Experimental models: Mouse Models** |  |  |
| Wild type C57BL/6J | Jackson | #000664 |
| VEGF<sup>lox/lox</sup> |  | Gerber et al.,<br>1999 |
| VEGF-GFP |  | Fukumura et al.,<br>1998 |
| Rosa-stop-floxed-EYFP | Jackson | Jackson #006148 |
| NestinCreER<sup>T2</sup> | Jackson | Jackson #016261 |
| Nestin-GFP | Jackson | Jackson #033927 |
| **Experimental models: Lentiviral Vectors** |  |  |
| mCherry | Kirby et al., 2015 |  |
| mCherry Cre | Kirby et al., 2015 | Addgene no.<br>27546 |
| GFP+ scramble shRNA | Mosher et al., 2012 |  |
| GFP+ *Vegfa* shRNA | Mosher et al., 2012 |  |
| CRISPRi non target | Addgene | TBD |
| CRISPRi αVEGFR2 | Addgene | TBD |
| **Software and algorithms** |  |  |
| ImageJ | schneider | https://imagej.nih.gov/ij/ |
| Zen | Zeiss | https://www.zeiss.com/microscopy/us/products/microscope-software/zen.html |
| Cell Ranger Cloud Analysis 7.0.1 | 10x Genomics | Zheng et al., 2017 |
| Seurat 4.1.1 | Satija Lab | Hao et al., 2021 |
| Cytoscape 3.9.1 |  | Shannon et al.,<br>2003 |
| ClueGO |  | Bindea et al.,<br>2009 |
| FlowJo 10.8.1 | BD Life Sciences |  |
| Image Studio | LI-COR |  |
| Prism | GfaphPad | https://www.graphpad.com/scientific-software/prism/ |
| **Other** |  |  |
| Batiuk et al., 2020 |  | doi:<br>10.1038/s41467-019-14198-8 |
| Hochgerner et al., 2018 |  | doi:<br>10.1038/s41593-017-0056-2 |
| Walker et al., 2020 |  | doi:<br>10.1186/s12967-020-02247-6 |
| Denninger et al., 2020 |  | doi:<br>10.1016/j.brainres.2020.146717 |
| Berg et al., 2019 |  | doi:<br>10.1016/j.cell.2019.02.010 |
| Adusumilli et al., 2021 |  | doi:<br>10.1016/j.stem.2020.10.019 |

## Supplementary figures

**Supplementary Figure 1:**
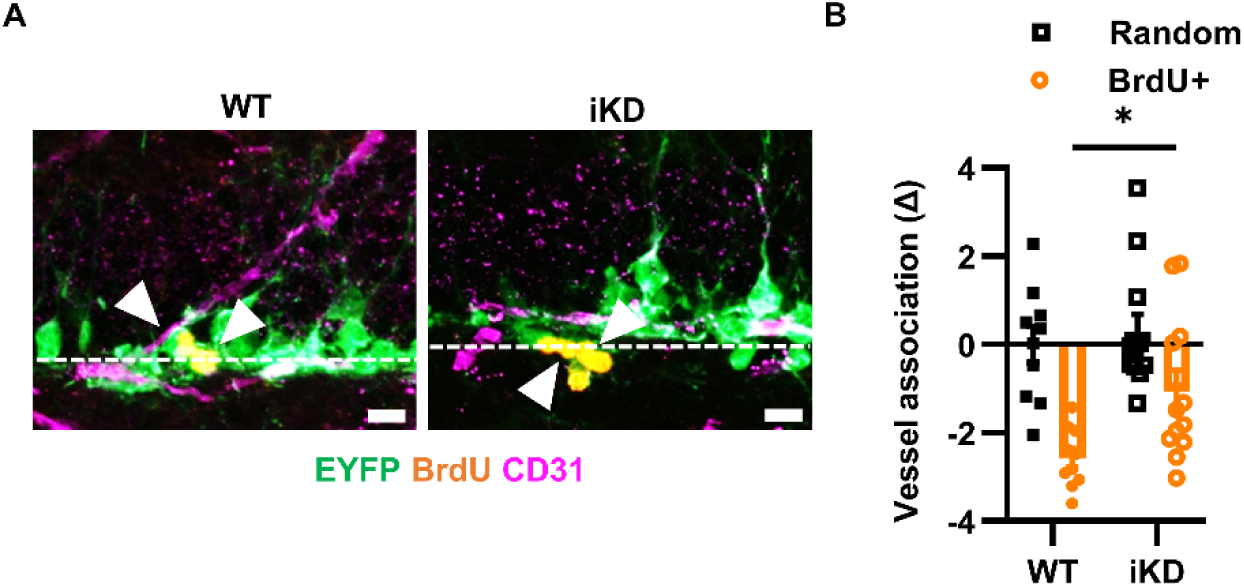
Vessel association of BrdU+ cells after VEGF iKD, related to **Figure 2**. A) Representative immunofluorescent images of BrdU+/EYFP+ cells and CD31+ endothelia 21d after NSPC-VEGF knockdown. Chevrons indicate BrdU+/EYFP+ cells. B) Vessel association measurement in WT or iKD BrdU+/EYFP+ cells compared to random point in SGZ. Mean ± SEM plus individual mice shown. N = 10 WT, 12 iKD. Scale bars represent 10 µm. *p < 0.05. Dashed line indicates lower border of granule cell layer.

**Supplementary Figure 2:**
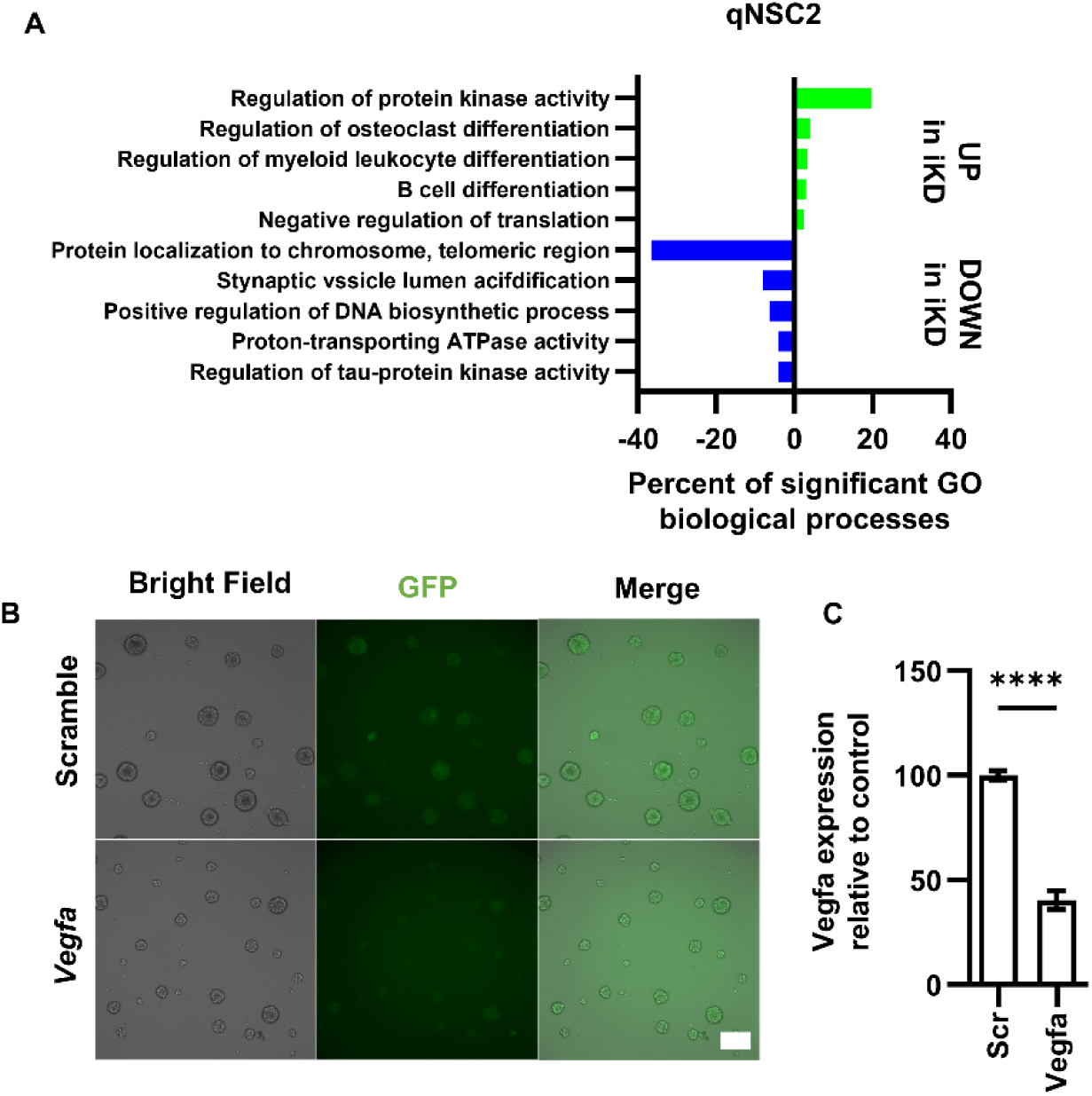
qNSC2 GO terms and *Vegfa* shRNA confirmation in vitro, related to Figures 4 and 5. A) Top 5 GO biological process clusters that were upregulated or downregulated in VEGF iKD qNSC2 cluster. B) Representative images of live cultured WT NSC spheres (bright field) after Scramble or *Vegfa* shRNA lentiviral infection (GFP+). C) Relative VEGF protein in conditioned media of WT NSCs after Scramble or *Vegfa* shRNA infection. N = 3/grp/exp, 2 exps; mean ± SEM. Scale bars represent 100µm. ****p < 0.0001.

**Supplementary Figure 3:**
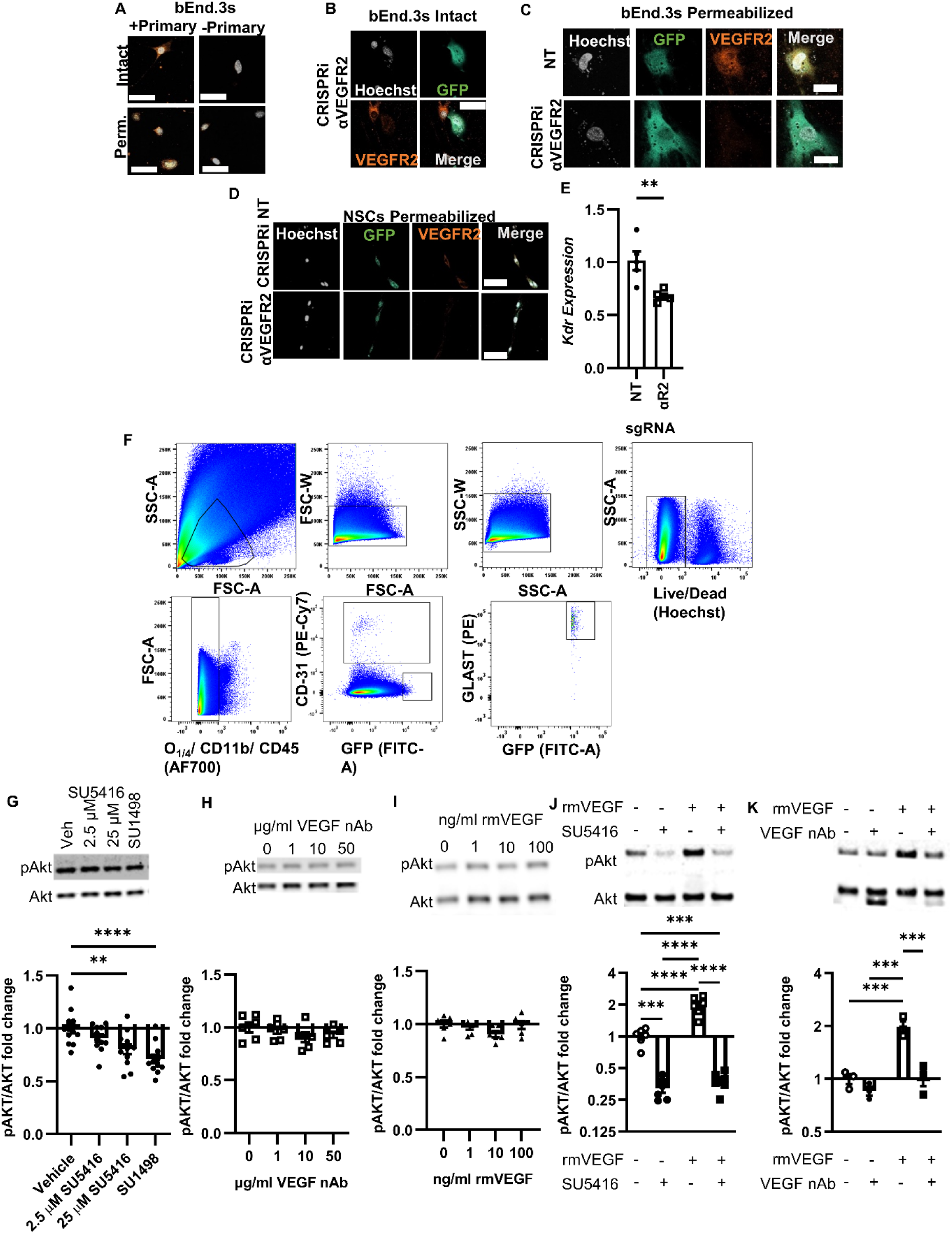
VEGFR2 antibody confirmation and further data on NSC insensitivity to extracellular VEGF, related to Figure 6. A) Representative VEGFR2 immunoreactivity in permeabilized and intact cultured bEnd.3s in VEGFR2 primary and no primary conditions. C) Representative images of VEGFR2 immunoreactivity in intact (B) or permeabilized (C) cultured bEnd.3s following transfection with CRISPRi NT or αVEGFR2. D) Representative images of VEGFR2 immunoreactivity in permeabilized cultured NSCs following infection with CRISPRi NT or αVEGFR2. E) *Kdr* mRNA expression in bEnd3.s after transfection with plasmids expressing an sgRNA targeting VEGFR2. N = 3 wells/grp/exp, 1 exp; mean ± SEM. F) Representative flow cytometry profiles for endothelial and NSC subsets in Fig 6G,H. G) Representative western blot of Akt phosphorylation in cultured DG NSCs after treatment with SU5416 or SU1498 (top). Fold change in Akt phosphorylation after SU5416 or SU1498 treatment in cultured NSCs (bottom). N = 2-3/grp/exp, 3 exps; mean ± SEM. H) Representative western blot of Akt phosphorylation in cultured DG NSCs after treatment with increasing dosages of VEGF nAb (top). Fold change in Akt phosphorylation after VEGF nAb treatment in cultured NSCs (bottom). N = 2-3/grp/exp, 3 exps; mean ± SEM. I) Representative western blot of Akt phosphorylation in cultured NSCs after treatment with recombinant mouse (rm)VEGF at increasing doses (top). Fold change in Akt phosphorylation after VEGF treatment in cultured NSCs (bottom). N = 3/grp/exp, 2 exps; mean ± SEM. J) Representative western blot of Akt phosphorylation in cultured HUVECs after treatment with recombinant mouse VEGF and SU5416 (top). Fold change in Akt phosphorylation after VEGF and SU5416 treatment in cultured HUVECs (bottom). N = 2-3/grp/exp, 1-2 exps; mean ± SEM. K) Representative western blot of Akt phosphorylation in cultured HUVECs after treatment with recombinant VEGF and VEGF nAb (top). Fold change in Akt after VEGF and VEGF nAb treatment in cultured HUVECs (bottom). N = 3/grp/exp, 1 exp; mean ± SEM. Scale bars = 50 µM. **p < 0.01; ***p < 0.001; ****p < 0.0001.

**Supplementary Figure 4:**
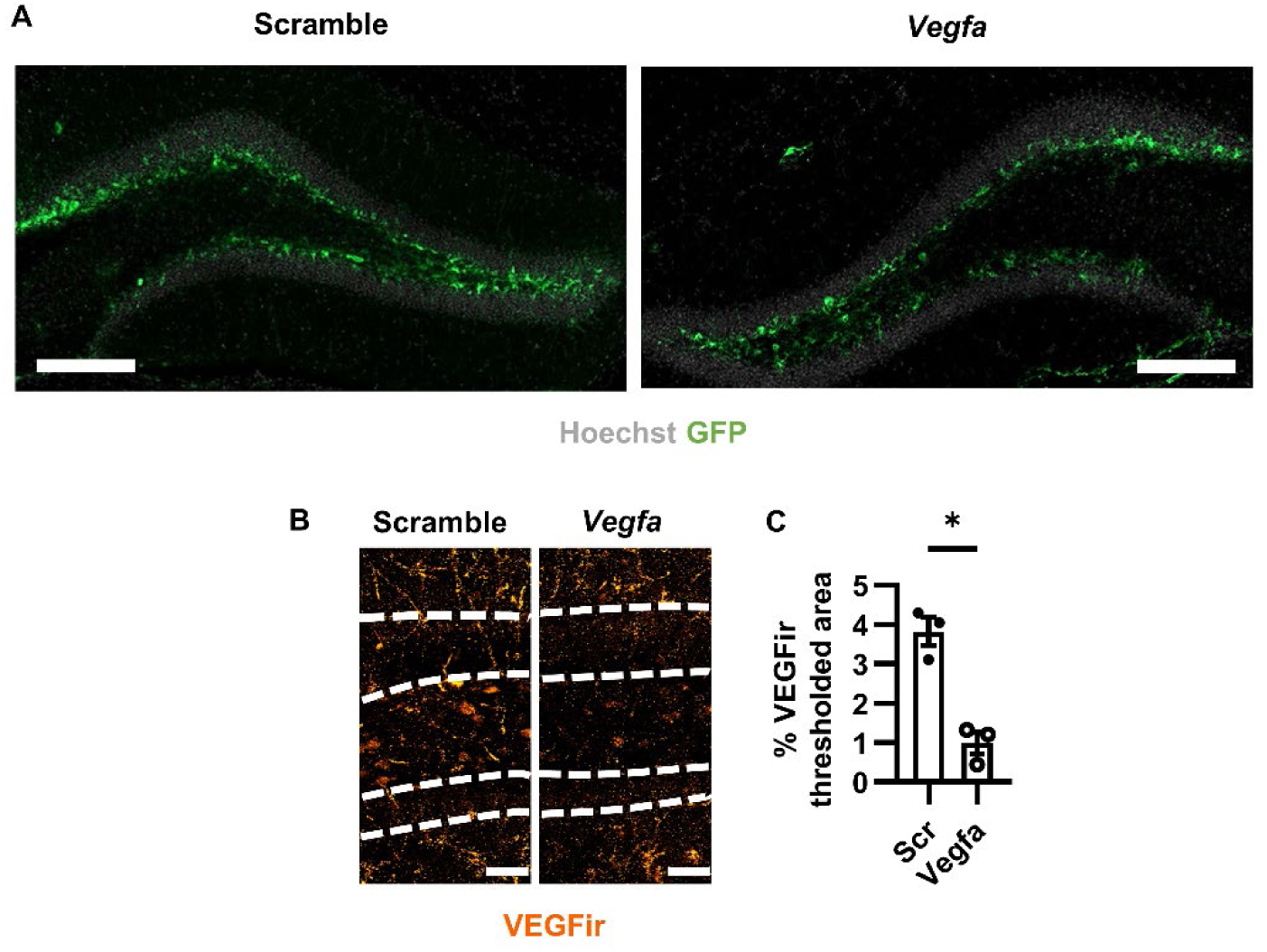
Lentiviral expression and VEGF immunolabeling, related to Figure 8. A) Representative images of Scramble or *Vegfa* shRNA infection (GFP+) in the DG (Hoechst shows cell nuclei) 21d after viral infusion. B) Representative VEGF immunoreactivity in the DG 21d after viral infusion. Dashed line indicates granule cell layer. C) Percent VEGF immunoreactive thresholded area in the DG 21d after viral infusion. N = 3 mice/grp. Mean ± SEM plus individual mice shown. Scale bars = A) 200 µM, B) 10 µM.

